# Modeling and Analysis of Hormone and Mitogenic Signal Integration in Prostate Cancer

**DOI:** 10.1101/058552

**Authors:** Katharine V. Rogers, Joseph A. Wayman, Ryan Tasseff, Caitlin Gee, Matthew P. DeLisa, Jeffrey D. Varner

**Affiliations:** School of Chemical and Biomolecular Engineering Cornell University, Ithaca NY 14853

**Keywords:** Prostate cancer, signal transduction, mathematical modeling

## Abstract

Prostate cancer is the most common cancer in men and the second leading cause of cancer related death in the United States. Androgens, such as testosterone, are required for androgen dependent prostate cancer (ADPC) growth. Androgen ablation in combination with radiation or chemotherapy remains the primary non-surgical treatment for ADPC. However, androgen ablation typically fails to permanently arrest cancer progression, often resulting in castration resistant prostate cancer (CRPC). In this study, we analyzed a population of mathematical models that described the integration of androgen and mitogenic signaling in androgen dependent and independent prostate cancer. An ensemble of model parameters was estimated from 43 studies of signaling in androgen dependent and resistant LNCaP cell lines. The model population was then validated by comparing simulations with an additional 33 data sets from LNCaP cell lines and clinical trials. Analysis of the model population suggested that simultaneously targeting the PI3K and MAPK pathways in addition to anti-androgen therapies could be an effective treatment for CRPC. We tested this hypothesis in both ADPC LNCaP cell lines and LNCaP derived CRPC C4-2 cells using three inhibitors: the androgen receptor inhibitor MDV3100 (enzalutamide), the Raf kinase inhibitor sorafenib, and the PI3K inhibitor LY294002. Consistent with model predictions, cell viability decreased at 72 hrs in the dual and triple inhibition cases in both the LNCaP and C4-2 cell lines, compared to treatment with any single inhibitor. Taken together, this study suggested that crosstalk between the androgen and mitogenic signaling axes led to robustness of CRPC to any single inhibitor. Model analysis predicted potentially efficacious target combinations which were confirmed by experimental studies in multiple cell lines, thereby illustrating the potentially important role that mathematical modeling can play in cancer.

## Introduction

Prostate cancer (PCa) is the most commonly diagnosed cancer and the second leading cause of cancer-related death in men in the United States [78]. Initially, PCa cells depend upon the activation of cytosolic androgen receptors (AR) by androgen hormones, such as testosterone, for survival and growth. Thus, androgen ablation in combination with radiation or chemotherapy remains the primary non-surgical treatment for androgen dependent prostate cancer (ADPC) [42]. However, androgen ablation typically fails to permanently arrest cancer progression as malfunctioning cells eventually lose androgen sensitivity and proliferate without hormone. The loss of androgen sensitivity results in castration resistant prostate cancer (CRPC), a phenotype closely linked with metastasis and reduced survival [34]. Currently, there are six approved treatments demonstrating a survival advantage in patients with metastatic CRPC, each target different aspects of the disease [72]. The taxane family members docetaxel and cabazitaxel interact with microtubule stability [19, 86], while abiraterone [72] and enzalutamide [74] interfere with androgen signaling by blocking androgen formation and nuclear translocation, respectively. Other treatments are not specific to PCa. For example, sipuleucel-T, a first generation cancer vaccine [44], or radium-223, an alpha emitter which targets bone metastasis [64], are both approved to treat CRPC. Unfortunately, regardless of the therapeutic approach, the survival advantage of these treatments is typically only a few months. Thus, understanding the molecular basis of the loss of androgen sensitivity in CRPC is an important step for the development of effective therapeutic strategies.

Androgen-induced proliferation and survival depends upon coordinated signal transduction and gene expression events. AR is a member of the nuclear hormone receptor superfamily, which includes other important receptors such as progesterone receptor (PR) and estrogen receptor (ER) [2]. Nuclear hormone receptors act as ligand dependent transcription factors interacting with specific DNA sequences on target genes as either monomers, heterodimers, or homodimers; AR, PR, and ER act as homodimers. For AR, these specific DNA sequences are called androgen response elements (ARE) [58]. In the absence of androgen, AR is predominately found in the cytoplasm bound to chaperones such as heat shock protein (HSP) [70]. Androgens, either testosterone or testosterone metabolites such as 5a-dihydrotestosterone (DHT), interact with cytosolic AR, promoting the dissociation of AR from HSP [69] and its subsequent dimerization, phosphorylation and translocation to the nucleus [4]. Activated nuclear AR drives a gene expression program broadly referred to as androgen action, that promotes both proliferation and survival. In addition to many genes including itself, activated nuclear AR promotes the expression and secretion of prostate specific antigen (PSA), arguably the best known PCa biomarker [22], although its prognostic ability is controversial [3, 40, 61].

Androgen dependent (AD) prostate cells become castration resistant (CR) through several possible mechanisms, including constitutively amplified AR expression, or altered AR sensitivity to testosterone or other non-androgenic molecules [22]. In this study, we focused on a third possible mechanism, the aberrant activation of AR by kinase signaling cascades, sometimes called the outlaw pathway. Outlaw activation can be driven by over-or constitutively activated receptor tyrosine kinases (RTKs), a common pathology in many cancer types including PCa [16, 79]. RTKs stimulate downstream kinases, including the AKT and mitogen-activated protein kinase (MAPK) pathways, which promote AR phosphorylation and dimerization in the absence of androgen [16, 96]. Interestingly, among the few genes activated AR represses is cellular prostatic acid phosphatase (cPAcP), itself a key negative regulatory of RTK activation [90]. Thus, in CRPC the androgen program is initiated without the corresponding extracellular hormone cue, potentially from crosstalk between growth factor and hormone receptor pathways. In turn, aberrant androgen action downregulates negative regulators of its own activation thereby forming a reinforcing positive feedback loop.

In this study, we analyzed a population of mathematical models that described the integration of androgen and mitogenic signaling in androgen dependent and independent prostate cancer. The model architecture was a significant advance over our previous prostate signaling model [87]. We added the regulated expression of ten additional proteins, including the cell cycle restriction point protein cyclin D, and included the regulation of AR action by cyclin D1a and the E2F transcription factor. We estimated model parameters using multiobjective optimization in combination with dynamic and steady-state data sets generated in AD, intermediate and CR PCa cell lines. We identified a population of models which described both AD and CR data sets using a single model structure. An ensemble of model parameters was estimated from 43 studies of signaling in androgen dependent and resistant LNCaP cell lines. The model population was then validated by comparing simulations with an additional 33 data sets from LNCaP cell lines and clinical trials. Analysis of the model population suggested that simultaneously targeting the PI3K and MAPK pathways in addition to anti-androgen therapies could be an effective treatment for CRPC. We tested this hypothesis in both ADPC LNCaP cell lines and LNCaP derived CRPC C4-2 cells using three inhibitors: the androgen receptor inhibitor MDV3100 (enzalutamide), the Raf kinase inhibitor sorafenib, and the PI3K inhibitor LY294002. Consistent with model predictions, cell viability decreased at 72 hrs in the dual and triple inhibition cases in both the LNCaP and C4-2 cell lines, compared to treatment with any single inhibitor. Taken together, this study suggested that crosstalk between the androgen and mitogenic signaling axes led to robustness of CRPC to any single inhibitor. Model analysis predicted efficacious target combinations which were confirmed by experimental studies in multiple cell lines, thereby illustrating the potentially important role that mathematical modeling can play in cancer.

## Results

**Estimating a population of prostate signaling models**. We modeled the integration of growth factor and hormone signaling pathways in AD and CR LNCaP cells (Fig. 1). The signaling architecture was curated from over 80 primary literature sources in combination with biological databases. We modeled both protein-protein interactions, and gene expression reactions involved in hormone and mitogenic signaling (Materials and Methods). The model equations were formulated as a system of ordinary differential equations (ODEs), where biochemical reaction rates were modeled using mass action kinetics. We estimated an ensemble of possible parameter sets using the Pareto Optimal Ensemble Techniques (POETs) algorithm [81]. POETs uses a combination of simulated annealing and local optimization techniques coupled with Pareto optimality-based ranking to simultaneously optimize multiple objective functions. Starting from an initial best fit set, we estimated the unknown model parameters using 43 *in vitro* data sets taken from six AD, intermediate and CR LNCaP cell lines (Table T1). Each of the training data sets was a separate objective in the multiobjective optimization calculation. The training data were steady-state or dynamic immunoblots from which we extracted relative species abundance using their optical density profiles. POETs generated well over a million possible parameter sets, from which we selected the top N = 5000 sets for further analysis. The coefficient of variation (CV) of the parameter ensemble spanned 0.59 – 5.8, with 33% of the parameters having a CV of less than one (Fig. S1). As a control, we also performed simulations for R = 100 random parameter sets to compare against the parameters estimated by POETs.

**Fig. 1:**
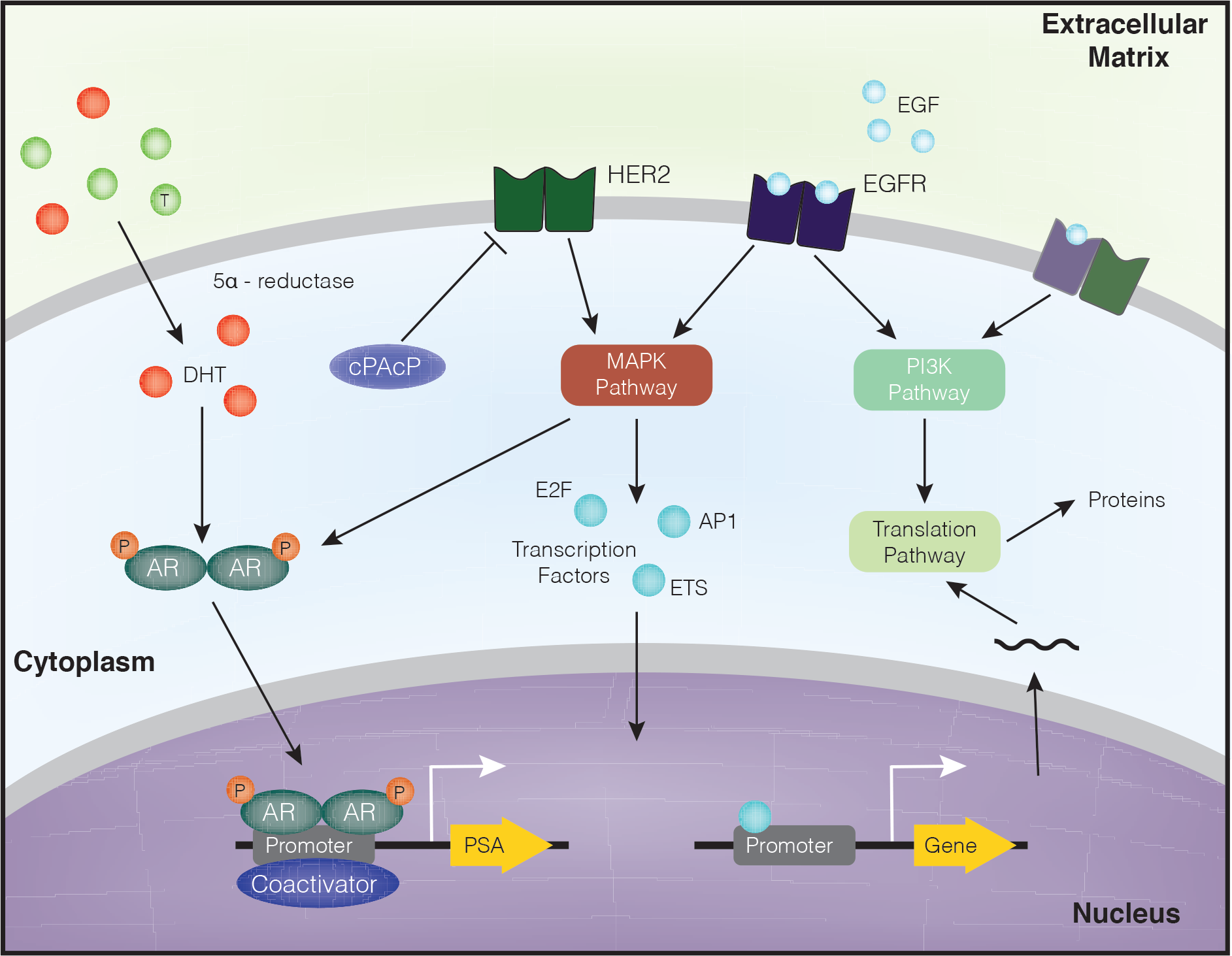
Schematic overview of the prostate signaling network. The model describes hormone and growth factor induced expression of several proteins, including PSA. In the absence of outside hormones/growth factors, overactive HER2 can stimulate the MAPK and AKT pathways. AR can be activated directly by the MAPK pathway.

**Table T1:**
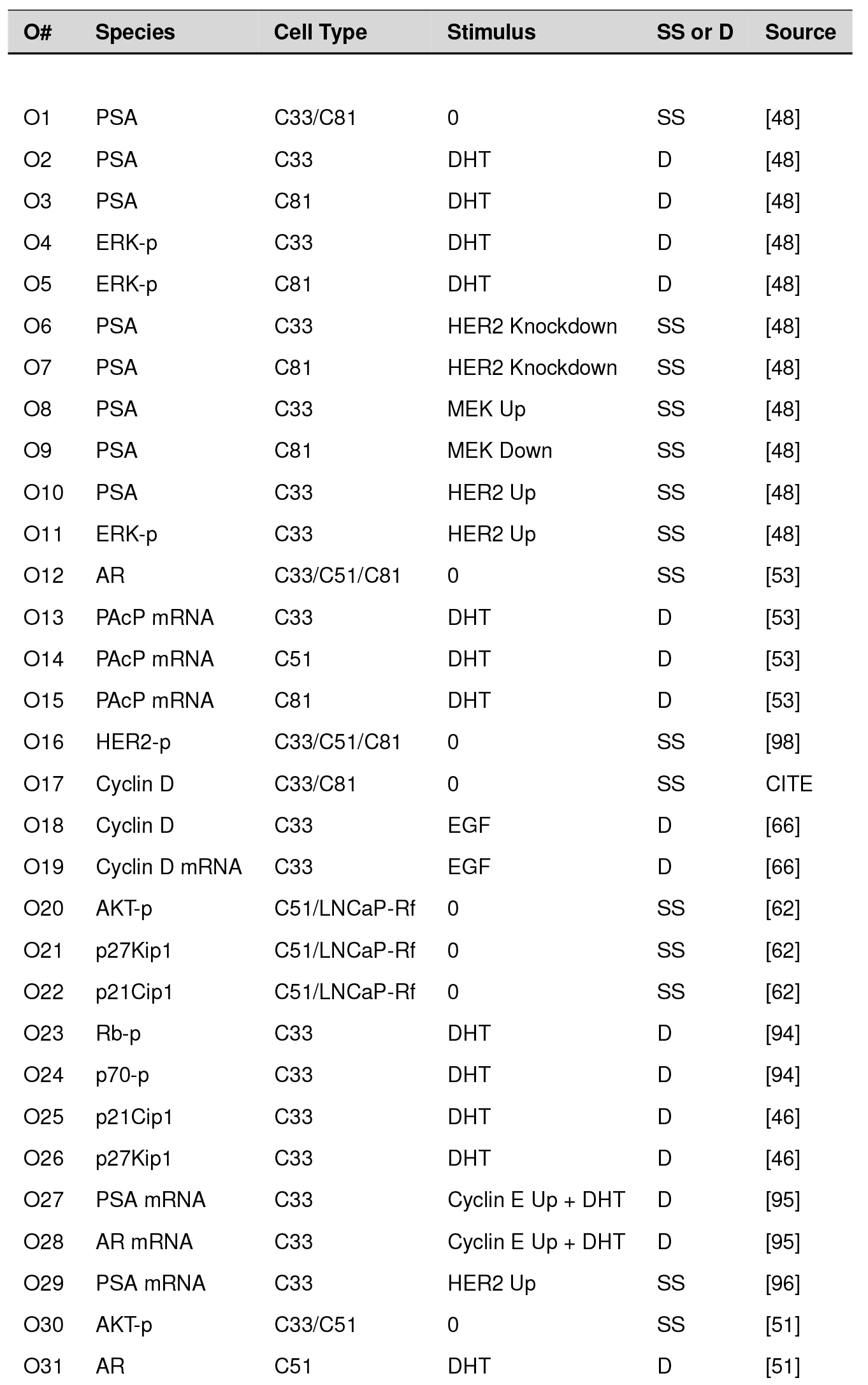
Objective function list along with species measured, stimulus, cell-type, steady state (SS) vs dynamic (D) and the corresponding literature reference.

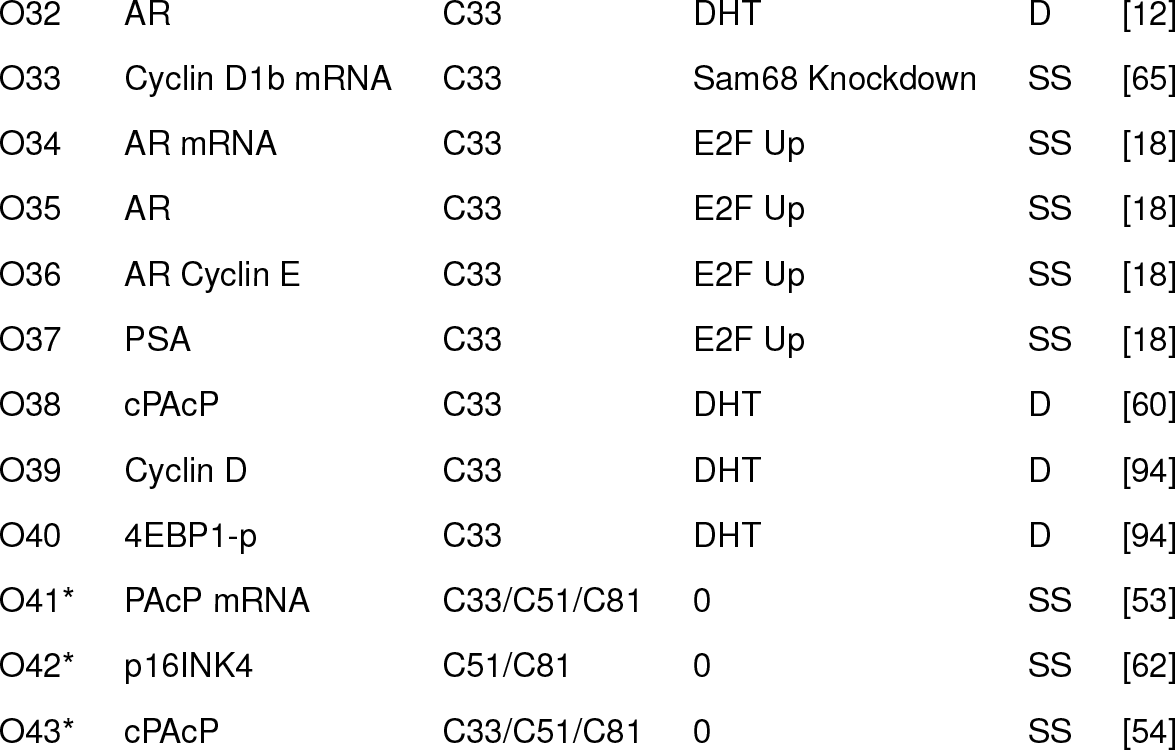

The population of signaling models recapitulated training data in both AD and CR cell lines with two experimentally mandated parameter changes (Fig. 2 and Fig. 3). Data from the LNCaP clones C-33 (dependent), C-51 (intermediate), and C-81 (resistant) [41, 43, 53] along with the CR LNCaP cell lines LNCaP-Rf [62], LNCaP-AI [11] and LNAI [26] were used for model identification. To simulate the effective difference between LNCaP cell lines, the parameter controlling the maximum rate of PAcP gene expression was scaled by 0.1 and 0.5, respectively, for the C-81 and C-51 cell-lines compared to C-33. This modification was based upon steady-state PAcP data from the three LNCaP clones [48]. Similarly, the expression of p16INK4 was adjusted in accordance with the study of Lu *et al.* [57]. These two parameters were the only adjustable parameter differences between AD and CR cells. To simulate an increased mTOR activation in the presence of a DHT stimulus, we added a first order activation term for mTOR activation with a DHT stimulus. Androgens increase the expression of proteins involved in cellular metabolism, leading to increased mTOR activation [94]. Conservatively, the model ensemble described approximately 85% of the training objectives (Fig. 2A), while only 20% of the training objectives were captured with the random parameter control (Fig. 2B). Thus, POETs identified a population of models that described the training data significantly better than a random parameter control.

**Fig. 2:**
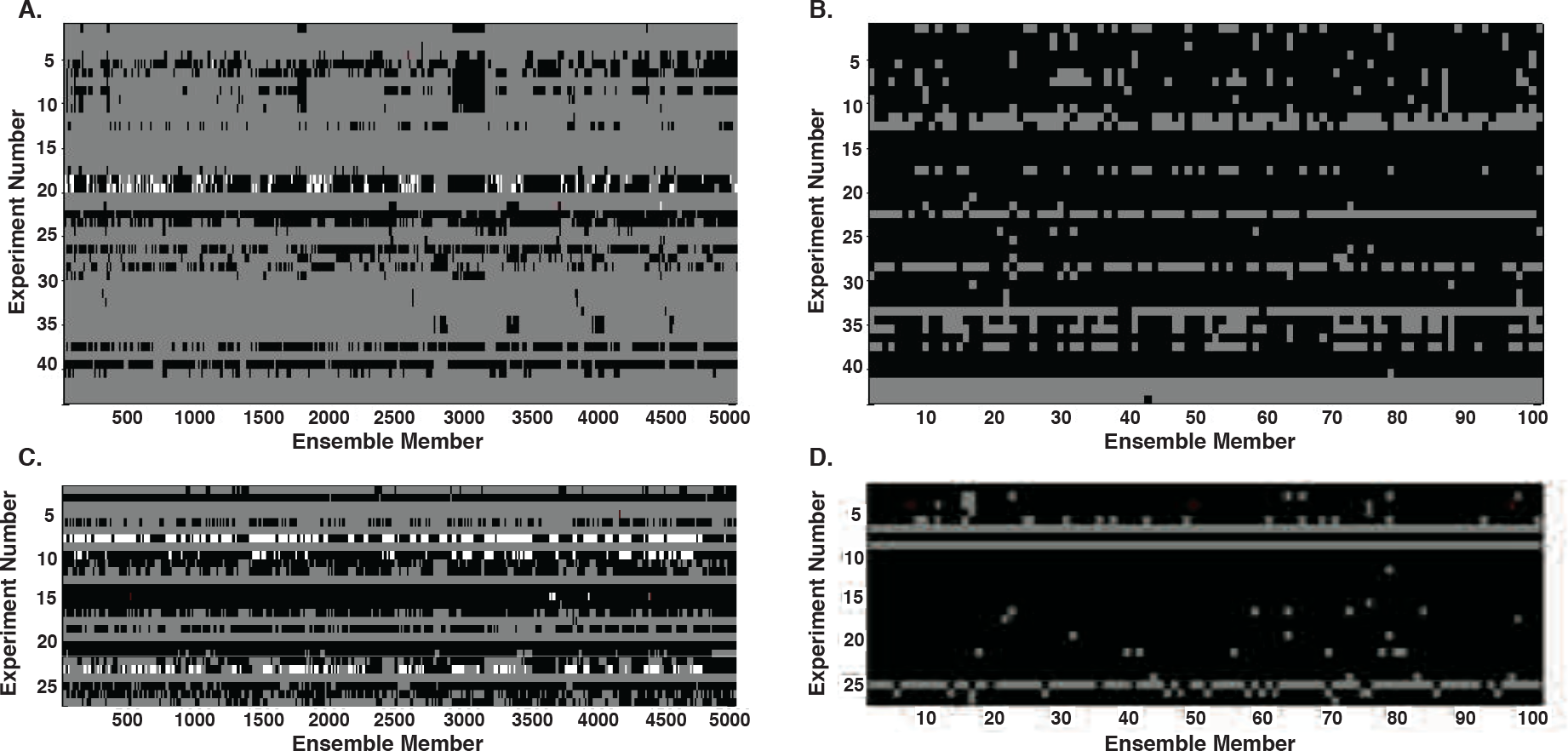
Simulation results versus experimental results for training and validation data. Experiment numbers 1 through 43 were used for training, while experiments 44 through 72 were validation. Gray means the ensemble member qualitatively fit experimental data in both models. White means the the ensemble member only fit the data using the new model that included HER2 heterodimerization. Red means the ensemble member fit using only the old model. Black corresponds to an incorrect cellular response in both models. A., C. Training and validation results, respectively, for entire ensemble population using both the original model and an updated model including HER2 heterodimerization (N = 5000). B., D. Simulation results for training and validation of a random set of 100 members using both models.

The population of models captured the crosstalk between RTK activation and androgen action (Fig. 3). The model described DHT-induced PSA expression (PSA is an AR-inducible gene) in both C-33 (Fig. 3A) and C-81 (Fig. 3B) cells. Simulations with the HER2 inhibitor AG879 also recapitulated decreased PSA expression in C-81 cells in the absence of androgen, highlighting the crosstalk between RTK and androgen action (Fig. 3C); AR action also decreased the PAcP mRNA message, presumably leading to increased HER2 activity (Fig. 3D). The model also recapitulated the integration of androgen action with AR expression, G1/S cell cycle protein expression and AKT phosphorylation. For example, the model captured AR-induced AR expression following a DHT stimulus (Fig. 3H). Conversely, the transcription factor E2F inhibited AR transcription in LNCaP cells (Fig. 3I). Other cell cycle proteins were also integrated with androgen action. For example, the cyclin D1 abundance increased in CR compared to AD cells in the absence of androgen (Fig. 3E), while DHT induced p21Cip1 expression in C-33 cells (Fig. 3F). The level of phosphorylated AKT also increased in higher passage number cells (Fig. 3G). Thus, the estimated model population recapitulated signaling and crosstalk behavior in both AD and CR LNCaP training data, above a random control. However, given the complexity of the model, it was unclear if the model ensemble could predict unseen data. To address this question, we fixed the model parameters and ran simulations of experimental data not used for model training.

**Fig. 3:**
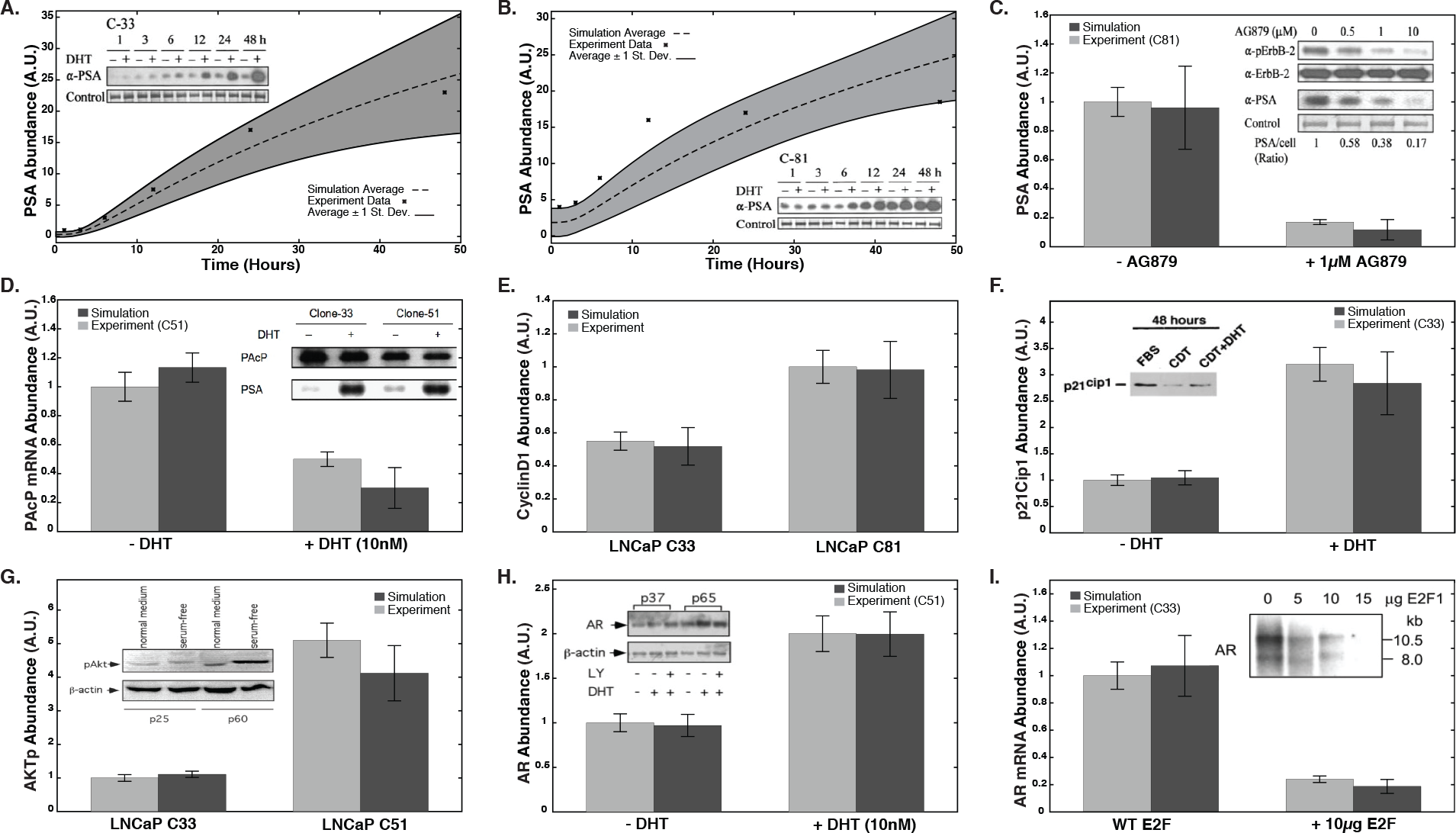
Ensemble performance against selected training objectives (N = 5000). A, B. Time course data for PSA concentration due to a stimulus of 10 nM DHT in LNCaP C33 cells and LNCaP C81 cells, respectively (O2, O3). C. PSA levels in the presence and absence of a HER2 inhibitor (LNCaP C81 cells, O7). D. PAcP mRNA levels at 72 hours in the presence and absence of DHT (LNCaP C51 cells, O14). E. Steady-state cyclin D levels in LNCaP C33 vs. C81 (O17). F. p21Cip1 levels at 48 hrs in the presence and absence of DHT (LNCaP C33, O25). G. Steady-state AKT phosphorylation levels in LNCaP C33 vs. C51 (O30). H. AR levels at 24 hours in the presence and absence of DHT (LNCaP C51, O31). I. AR mRNA levels in the presence and absence of E2F over expression (LNCaP C33, O34). Error bars denote plus and minus one standard deviation from the mean.

**Validation simulations revealed missing network structure**. The model was validated against 29 *in vitro* and four in vivo clinical studies (Table T2). For 15 of the 29 cases, the ensemble was qualitatively consistent with the experimental data (Fig. 2C). However, for the random parameter control, only 7 of the 29 cases were satisfied (Fig. 2D). We correctly predicted positive feedback between HER2 auto-activation and androgen action (Fig. 4A and Fig. 4B). We also captured the dose-dependence of AR abundance on DHT (Fig. 4C). In addition to the cell line studies, we simulated the outcome of enzalutamide, lapatinib, and sorafenib clinical trials in AD and CRPC patients. The trial end points were the reduction in PSA expression relative to an untreated baseline. Enzalutamide acts on AR by inhibiting its nuclear translocation, DNA binding, and coactivator recruitment [74]. In the enzalutamide trial, 54% of the patients that received the drug showed a PSA decline of ≥ 50% while 25% showed a decline ≥ 90%. We simulated enzalutamide exposure by reducing the rate constants governing activated AR binding to nuclear importer, cyclin E, and CDK6 to 1% of their initial values. Consistent with the trial, 62% of ensemble members showed a ≥ 50% decline in PSA abundance, while 14% showed a ≥ 90% decline (Fig. 4G). Next, we simulated the response of our model population to lapatinib, an inhibitor of epidermal growth factor receptor (EGFR) and HER2 tyrosine kinase activity [55]. Two lapatinib drug trials were considered: one in which patients had CRPC and one in which patients had biochemically relapsed ADPC [55, 93]. In the CRPC lapatinib trial, 9.5% of the enrolled patients had a PSA response ≥ 47% [93], while our model ensemble showed 26% PSA response rate. Of the 35 patients enrolled in the ADPC lapatinib study, no PSA decreases was observed [55]; the model ensemble showed less than a 10% PSA response rate (data not shown). However, while no response to lapatinib was seen in ADPC clinical trials, *in vitro* AD LNCaP experiments showed decreased PSA expression in response to lapatinib, most notably with the addition of DHT [56]. Lastly, we simulated the response of CRPC patients to sorafenib, a kinase inhibitor with activity against Raf, vascular endothelial growth factor receptor (VEGFR), platelet-derived growth factor receptor (PDGFR), c-kit and c-Ret [17]. We considered only the effects of sorafenib on Raf, as the others were not included in the model. None of the 22 patients in the sorafenib study showed a PSA decline of > 50%. However, our simulations showed that approximately 55% of the ensemble members had a PSA decline of ≥ 50% (Fig. 4I). Taken together, the model ensemble predicted approximately 55% of the validation cases overall, but 75% of the clinical test cases. The failed clinical cases, and many of the failed training and validation cases, involved RTK activation, and in particular epidermal growth factor (EGF) signaling, suggesting the model was missing key biology.

Training and validation failures suggested the original signaling architecture was missing critical components related to EGF signaling. Several of the failed training and validation simulations involved the response of the network to EGF stimulation. For example, Chen *et al.* showed that HER2 phosphorylation increased within five minutes following EGF stimulation of LNCaP-AI cells [11]. However, we predicted no connection between HER2 phosphorylation and EGF stimulation on this short timescale (Fig. 4E). Interestingly, we initially neglected the heterodimerization of HER2 with other ErbB family members to simplify the model. Chen *et al.* suggested that HER2-EGFR heterodimerization was an important factor in EGF-driven activation of HER2 [11]. We tested this hypothesis by developing a new model that included HER2 and EGFR heterodimerization (all else held the same). We set the rate constants governing the assembly of HER2/EGFR heterodimers equal to EGFR homodimer assembly; all other parameters were unchanged. We felt this was a reasonable first approximation, as the affinity of HER2/EGFR heterodimerization and EGRF homodimerization is thought to be similar [38]. With the inclusion of HER2-EGFR heterodimerization, we qualitatively described EGF-induced HER2 activation, and more generally improved our training peformance for experiments that involved an EGF stimulus, e.g., cyclin D mRNA and protein abundance following an EGF stimulus in C-33 cells (Fig. 2A and C, white pixels and Fig. S2). This structural update to the model improved the training percentage to approximately 90%, and also highlighted an advantage of the ensemble modeling approach. Next, we analyzed the ensemble of models using both local and global techniques to estimate which parameters and processes were controlling system performance for AD and CR cells.

**Table T2:**
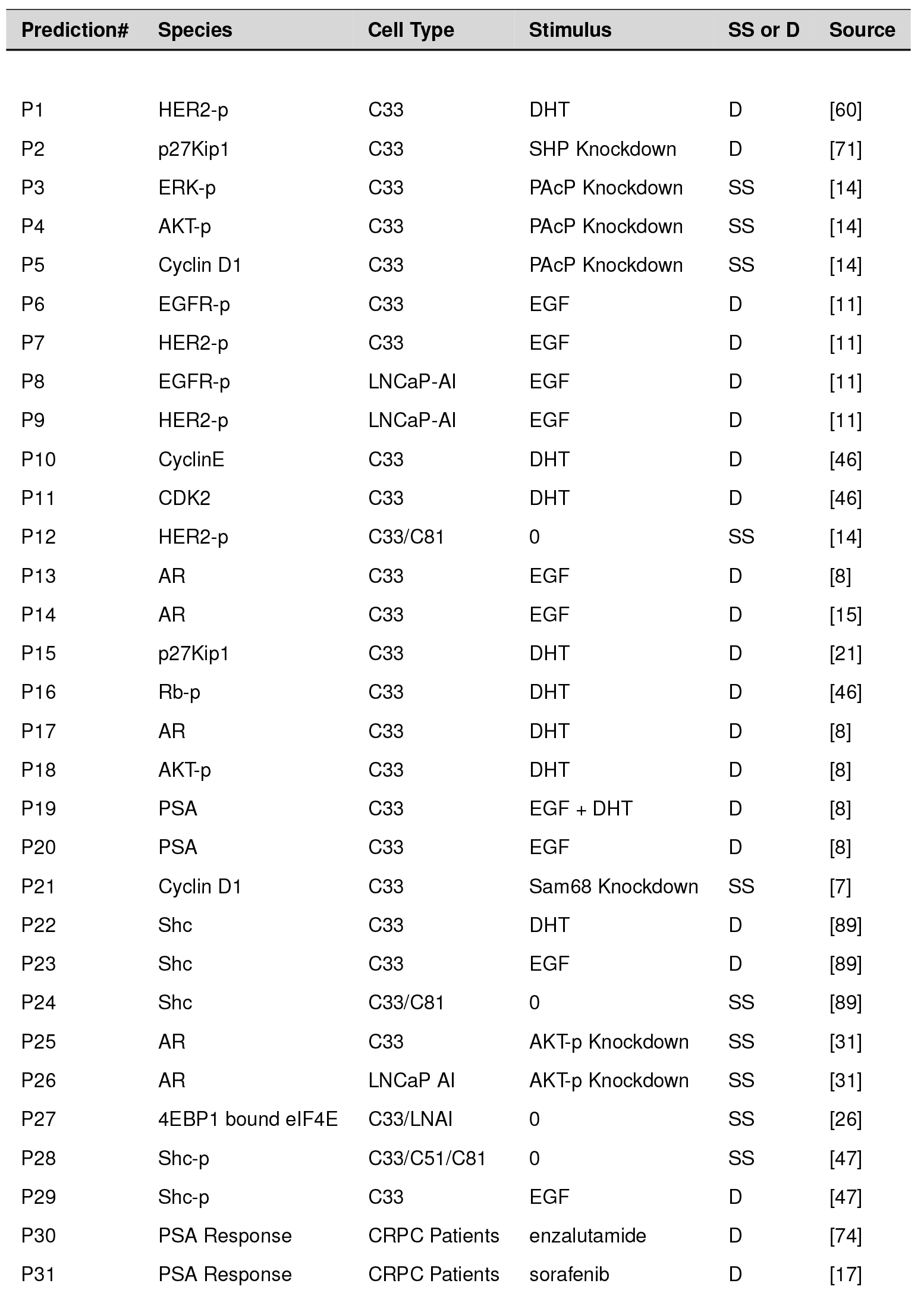
Blind Prediction list along with species measured, stimulus, cell-type, steady state (SS) vs dynamic (D) and the corresponding literature reference.

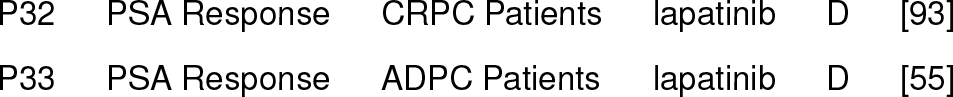

**Fig. 4:**
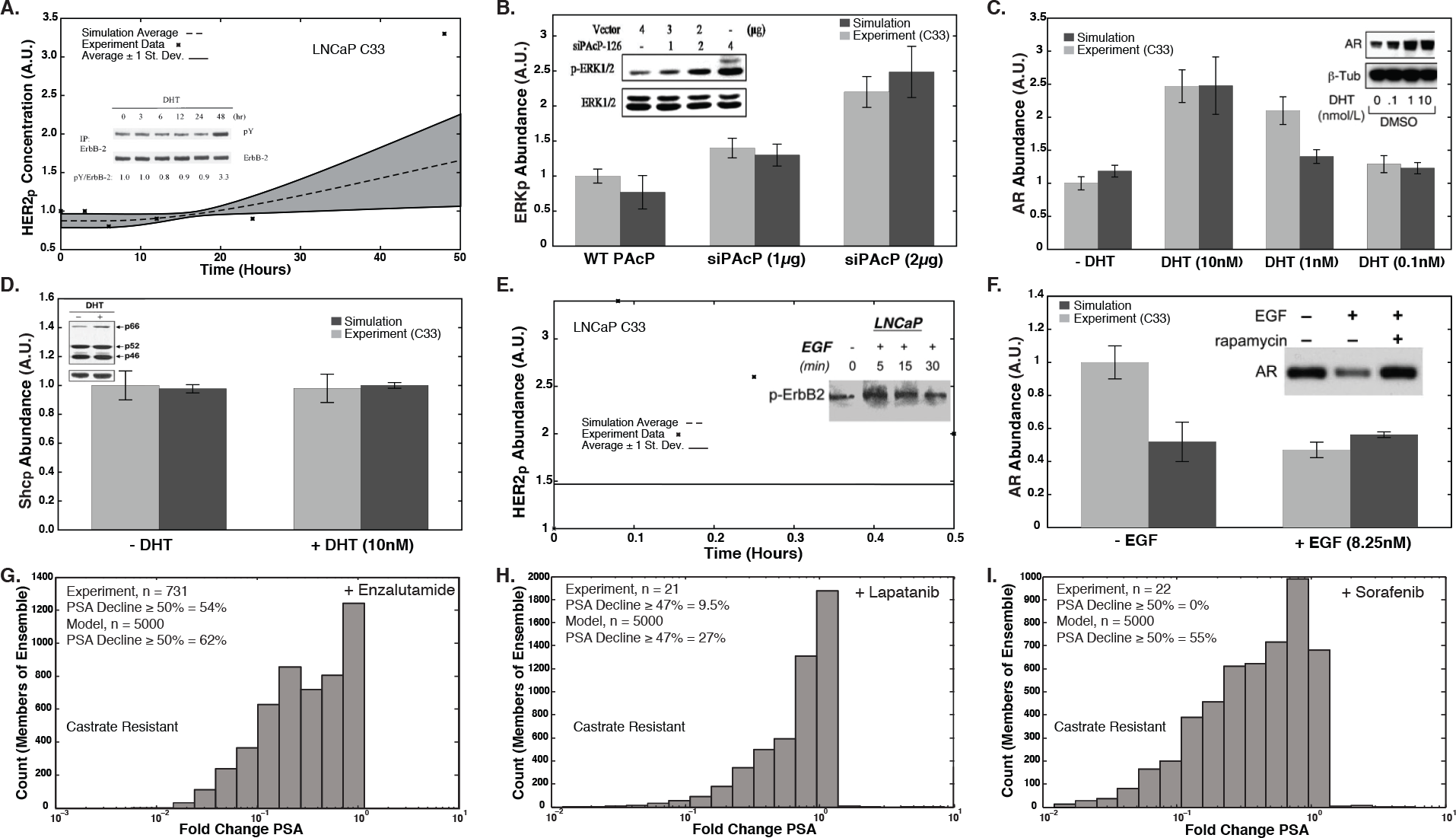
Blind model predictions for the ensemble (N = 5000). The model ensemble's predictive ability was assessed by comparing simulation versus experimental data not used for training. A. Time course data for HER2 phosphorylation due to a stimulus of 10 nM DHT (LNCaP C33, P1). B. ERK phosphorylation levels in the presence and absence of a PAcP inhibitor (LNCaP C33 cells, P3). C. AR levels at 24 hrs in varying levels of DHT (LNCaP C33, P17). D. Shc phosphorylation levels at 24 hrs in the presence and absence of DHT (LNCaP C33, P22). E. Time course data for HER2 phosphorylation due to a stimulus of 1.6 nM EGF (LNCaP C33, P7). F. AR levels in varying levels of EGF (LNCaP C33, P14). G, H, I. Fold change in PSA concentration due to drug stimulus: enzalutamide, lapatinib, and sorafenib. Error bars denote plus and minus one standard deviation from the mean.

**Sensitivity analysis identified differentially important network features**. Sensitivity analysis identified important signaling components in AD versus CR cells (Fig. 5). We calculated first order steady-state sensitivity coefficients under different stimuli for 500 parameter sets randomly selected from the ensemble. The sensitivity profile was similar for AD versus CR cells in the presence of DHT (Fig. 5B). The top 2% of sensitive species belonged to either the MAPK or PI3K pathways. In particular, activated Ras, Raf, phosphorylated MEK, PIP3 localized AKT, phosphorylated AKT, and PI3K were sensitive in both AD and CR cells. PAcP and p16INK4 along with E2F, cyclin E, and DHT-activated AR were more sensitive in AD cells. On the other hand, HER2 activation of PI3K, and AKT inhibition of Raf were more sensitive in CR cells. Taken together, in the presence of DHT, AD and CR cells shared a similar sensitivity profile with only a few differences. This suggested androgen had a strong influence on network performance even for CR cells. Next, we analyzed the ensemble of models in the absence of androgen.

**Fig. 5:**
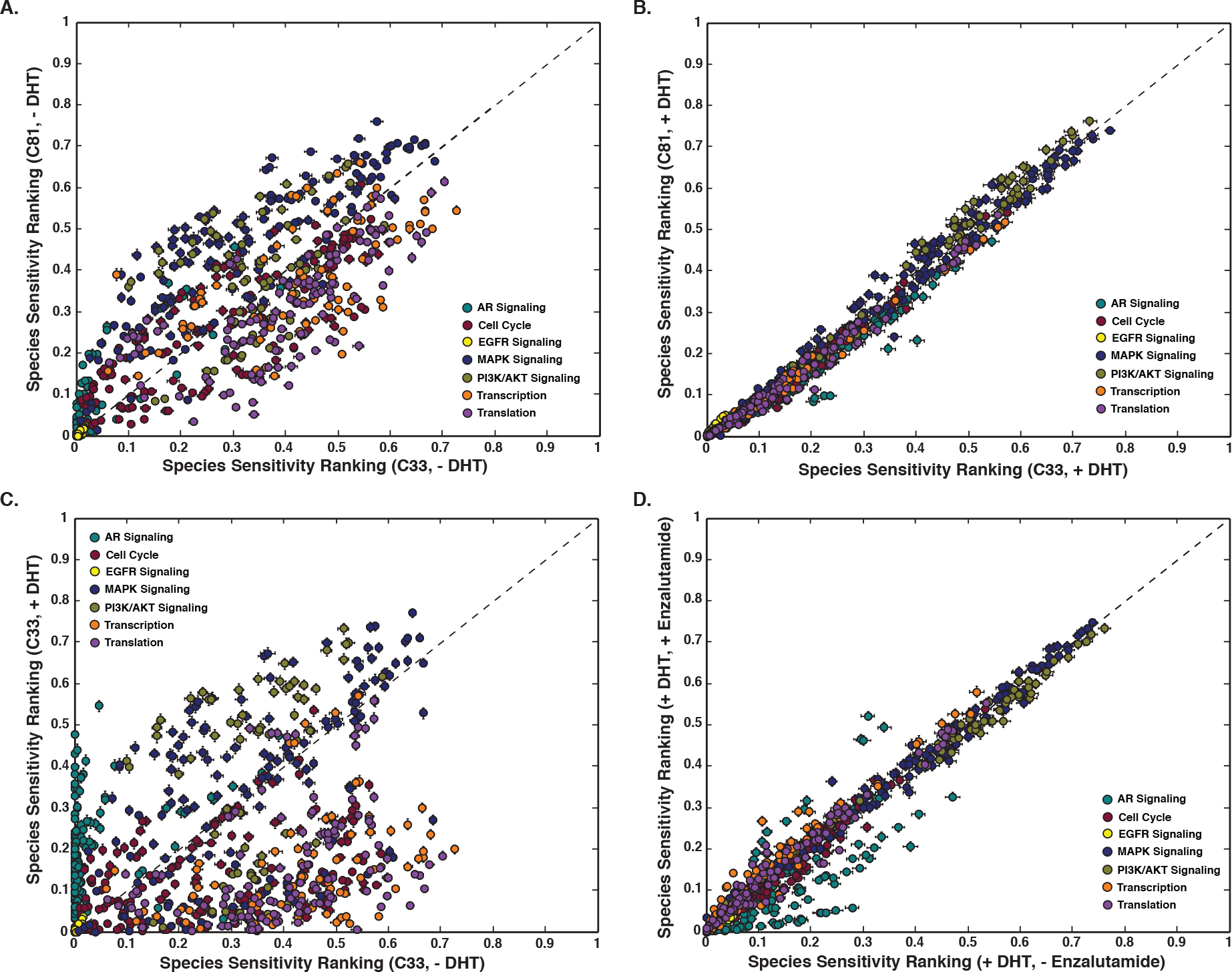
Sensitivity analysis of a population of prostate models (N = 500). Species with a low sensitivity are considered robust, while species with a high sensitivity ranking are considered fragile. A, B. Sensitivity ranking of network species in AD versus CR cells in the absence (presence) of DHT. C. Sensitivity ranking of network species in AD cells in the absence and presence of DHT. D. Sensitivity ranking of network species in CR cells in the presence and absence of enzalutamide with a DHT stimulus. Error bars denote standard error with N = 500.

The importance of signaling components varied with androgen dependence in the absence of DHT (Fig. 5A). There were 108 sensitivity shifts that were greater than one standard deviation above the mean shift. In CR cells, HER2 activation of ERK and PI3K was more sensitive, as was AR activation through the MAPK pathway. In general, the MAPK pathway was more sensitive, and sPAcP more robust in CR cells. This was expected, as outlaw pathway activity is elevated in castration resistant cells. On the other hand, infrastructure pathways encoding transcription and translation were more sensitive in AD cells. PSA and cyclin D1b (mRNA and mRNA complexes) were the only species involved in translation that were more robust in AD cells. This would suggest that the targeting of transcription or translation mechanisms in CR cells may be less effective than in AD cells. The transcription factor, E2F was more fragile in AD cells, while the transcription factors ETS and AP1 were more robust. The model included AP1 suppression of AR transcriptional activity (more sensitive in CR) [73], as well as inhibition of transcription of the AR gene by E2F (more sensitive in AD) [18]. Species in the PI3K pathway that were more fragile in AD cells included Rheb and TOR complexes. Interestingly, these species were included as the last step in the PI3K pathway prior to translation, with the phosphorylation of 4E-BP1 by TOR being considered the beginning of translation in this model. This again indicates that in the absence of DHT general translation is more fragile in AD cells.

Next we considered the importance of signaling components in the presence and absence of androgen for the same cell type. There were a total of 119 significant shifts between an androgen and a non-androgen environment in AD and CR cells (Fig. 5C and Fig. S3). Unsurprisingly, AR activation through DHT binding, with and without coactivators, in a DHT environment was more sensitive, as was AR inhibition of PAcP transcription (repressed by AR in the model). Species further upstream, such as HER2 activation of the MAPK and PI3K/AKT pathways, were also more sensitive in a DHT environment. This is most likely the result of the positive feedback between androgen action and HER2 activation in the model. Cell cycle species that were more fragile in the presence of DHT included complexes involving p21Cip1 and CDC25A. In a non-androgen environment, basal transcription and translation were more sensitive. Other sensitive species in the absence of DHT included Rb, E2F, Sam68, cyclin D1a complexes, MAPK phosphatases, and Rheb/TOR complexes. Notably, the value of the species sensitivity ranking shifts for basal transcription and translation in an androgen versus a non-androgen environment were higher in AD vs CR cells (Fig. 5C and Fig. S3). This again may indicate that in an androgen free environment in an AD cell, targeting of general translation and transcription may be beneficial, but may be less effective in a CR cell.

Lastly, we considered the sensitivity of CR cells in the presence of the AR inhibitor enzalutamide with and without DHT. The top 2% of sensitive species with and without enzalutamide were conserved in the presence of DHT (Fig. 5D). Species which were more sensitive with enzalutamide and DHT included cytosolic AR, cPAcP, and p21Cip1. As expected, nuclear AR was more robust in the presence of enzalutamide. Enzalutamide prevents translocation of AR to the nucleus causing levels of nuclear AR to decrease and cytosolic AR to increase. In CR cells, enzalutamide had no effect on the sensitivity of PI3K/AKT or MAPK species, many of which were included in the top 2% of sensitive species. Next, we looked at the effect of enzalutamide on CR cells without DHT (Fig. S3). Dimerized HER2, ERK, and PAcP were more sensitive in a non-androgen environment with enzalutamide. Species which were more robust in the non-androgen environment included, AR activated by DHT, AKT, p70, and AR bound to HSP. Taken together, the sensitivity results suggested that instead of inhibiting the AR pathway alone (enzalutamide), a combination approach targeting the PI3K or MAPK pathways in addition to AR could be more effective in treating CR cells. However, first-order sensitivity coefficients measure the result of infinitesimal changes to model parameters. Thus, they may not faithfully reflect the outcome of a finite perturbation to the network. To address this shortcoming, next we simulated the response of AD and CR cells to knockouts or amplification of network components.

**Robustness analysis confirmed the need for dual inhibition**. Robustness analysis was conducted to quantify the effects of amplifying or removing key model components in AD and CR cells. Gene expression parameters were altered by a factor of 10, 0.5, and 0 for knock-in, knock-down, or knock-out perturbations, respectively. We calculated the effect of these perturbations on the expression or activation of different protein markers, such as PSA, AR, cyclin D, activated p70 and phosphorylated AKT. In particular, we calculated the effect of knock-out perturbation in CR cells for seven cases: (1) Raf knockout, (2) PI3K knock-out, (3) AR knock-out, (4) Raf and PI3K knock-outs, (5) Raf and AR knock-outs, (6) PI3K and AR knock-outs, and (7) Raf, PI3K and AR knock-outs. Over the 500 models sampled, the greatest decrease in PSA expression occurred for cases involving AR knock-outs (Fig. 6A). On the other hand, the greatest decrease in activated p70 abundance occurred for the PI3K/AR and the triple Raf/PI3K/AR knock-out cases (Fig. 6B). The median and mean response for cyclin D expression was near zero for all knock-out cases (Fig.6C). However, there was significant variance over the population of models (both increased and decreased expression) in response to the perturbations. As expected, PI3K activity was required for AKT phosphorylation, while other knockouts had little influence on AKT phosphorylation (Fig. 6D). However, as was true with cyclin D, there was a subpopulation of models with increased AKT phosphorylation for cases involving RAF and RAF/AR knock-outs. These results support our case for a combination treatment approach, with PSA, activated p70, and AKT phosphorylation all decreasing in the PI3K/AR knock-out case as well as the RAF/PI3K/AR knock-out case.

**Fig. 6:**
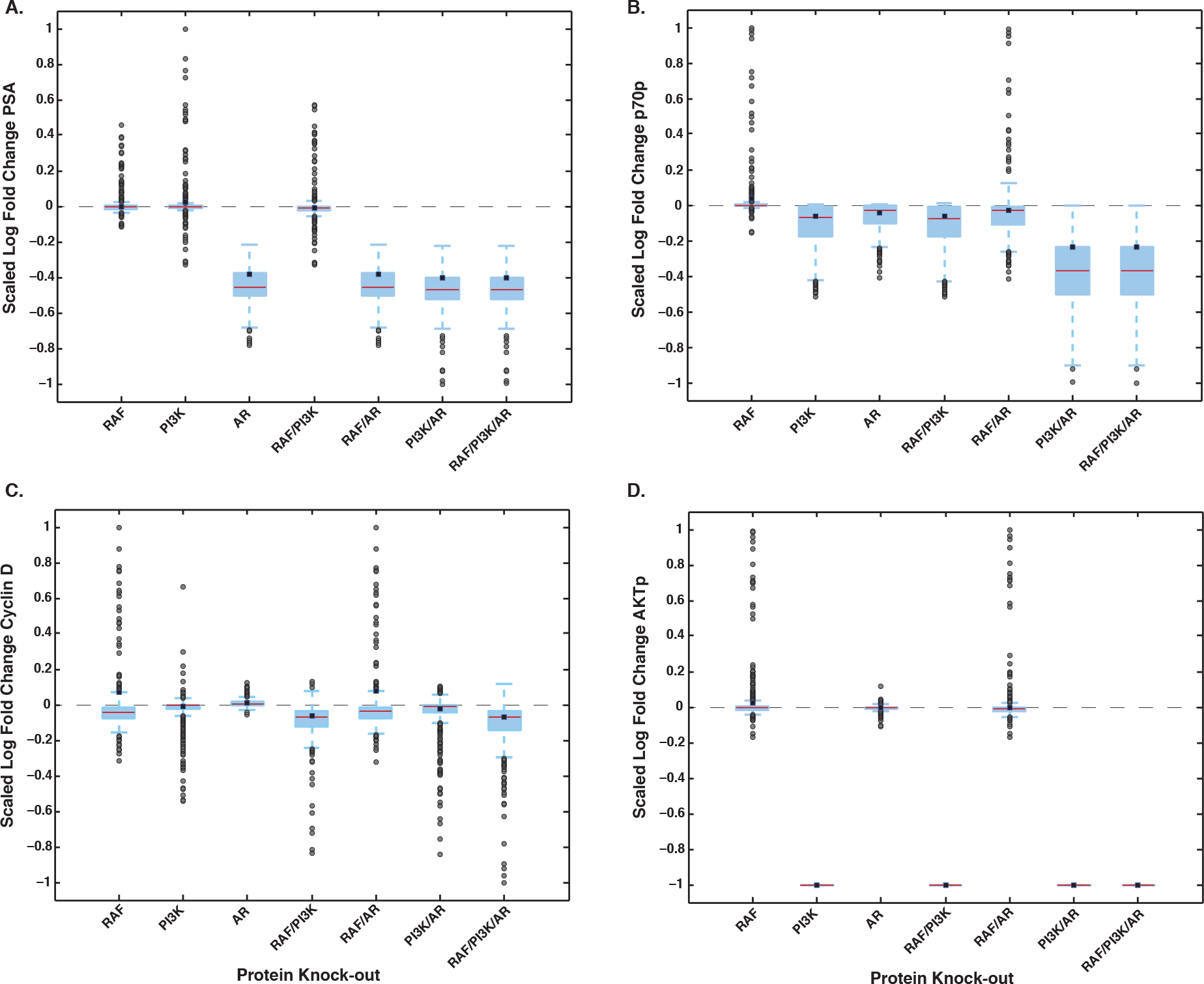
Robustness analysis of a population of CR prostate models with seven protein knock-out cases (N = 500). A scaled log fold change of greater than zero implies that the concentration of the protein increased with the knock-out, while a scaled log fold change of less than zero indicates that the concentration of protein decreased. A scaled log fold change equal to 0, shows no response due to the knock-out. A.,B.,C.,D. Log robustness of PSA, p70p, cyclin D, and AKTp versus protein knock-out. A CR LNCaP cell was assumed for all knock-out cases. The bottom and top of each box denotes 25th and 75th percentiles, while the red line indicates the median. The whiskers on the plot are plus and minus 1.5 the interquartile range (IQR) from the top and bottom values of the box, respectively. The grey dots denote outliers and the blue dots denote the mean.

Next we looked at single gene expression perturbations to understand the variance over the population of models. A knock-out of Raf, MEK or ERK showed an average overall increase in cyclin D levels in CR cells (Fig. S4). This was unexpected and we saw a similar increase in cyclin D due to the knock-in of Raf, MEK or ERK. We found that individual models showed different response to a Raf knock-out, in both cyclin D and PSA abundance. We saw three distinct regions: (1) increased PSA expression, (2) increased cyclin D expression, and (3) decrease in both PSA and cyclin D expression. Of the 500 models, 126 models had increased PSA expression, and 62 models had increased cyclin D expression due to the knock-out of Raf (Fig. 7). We explored the flux vectors of the outlying parameter sets to understand the mechanistic effect of Raf knock-out on PSA and cyclin D. Outlying parameter sets in region 1 displayed high activation of PI3K through HER2 signaling as well as high association of AP1 with AR. AP1 is known to bind and suppress AR transcriptional activity in LNCaP cells [73]. Knocking out Raf lowered AP1 levels and, therefore, freed AR for increased transcription of PSA. Models in region 2 also had high activation of PI3K through HER2, as well as higher association of E2F with Rb and cyclin D1a with AR. Cyclin D levels in region 2 increased due to an increase in E2F levels caused by the Raf knock-out. Models in region 3 had high association of mTOR. Interestingly, the drug sorafenib, a multi-kinase inhibitor that has activity against Raf, showed no measurable PSA decline in prostate cancer patients in clinical trials [17]. The robustness analysis showed that network perturbation can result in unexpected responses due to heterogeneity in signal transduction and gene expression processes.

**Fig. 7:**
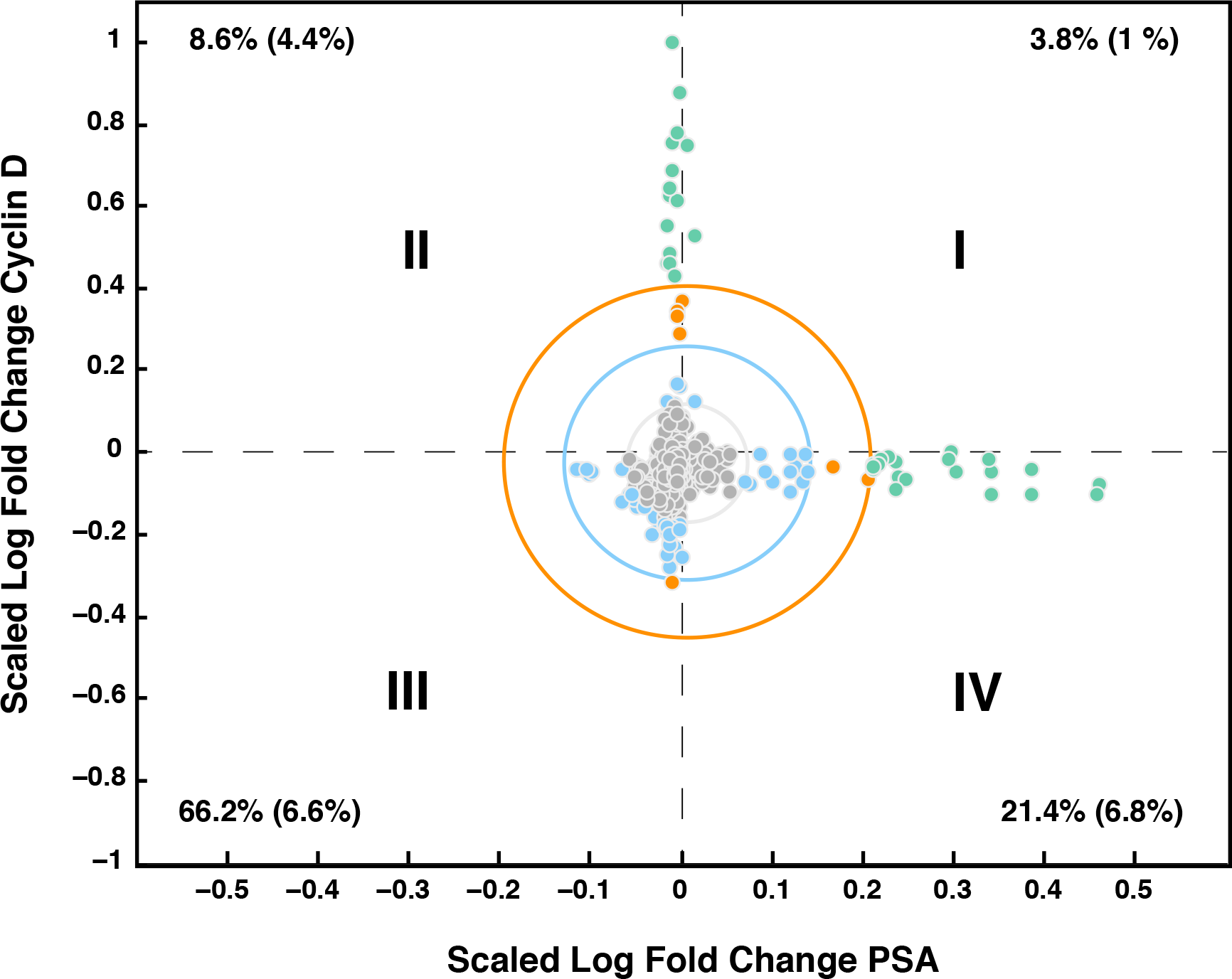
Robustness analysis of a population of CR prostate models with Raf knock-out (N = 500). A scaled log fold change of greater than zero implies that the concentration of the protein increased with the knockout of Raf, while a log fold change of less than zero indicates that the concentration of protein decreased. A log of fold change equal to 0, shows no response due to Raf knock-out. Three distinct regions emerge in Raf knock-out case: (1)PSA increases, (2)cyclin D concentration increases, and (3)PSA and cyclin D concentration decrease. The grey ellipse is centered at the mean values with an x-radius and y-radius of one standard deviation of the scaled log fold change of PSA values and cyclin D values, respectively. The blue ellipse denotes two standard deviations from the mean and the orange denotes three standard deviations from the mean. Values denote percentage of total parameter sets that fall in each quadrant, while values in parenthesis denote the percentage that fall at least one standard deviation from the mean.

**Experimental studies confirmed the effectiveness of dual and triple inhibition**. Sensitivity and robustness analysis, conducted over a subpopulation of prostate signaling models, suggested that simultaneously targeting the PI3K and MAPK pathways in addition to anti-androgen therapies could be an effective treatment for CRPC. To test this hypothesis, we measured the response of the well characterized ADPC cell line LNCaP as well a LNCaP derived CRPC cell line C4-2 to inhibitor and inhibitor combinations (Fig. 8). Three inhibitors were used: the AR inhibitor MDV3100 (enzalutamide), the Raf kinase inhibitor sorafenib, and the PI3K inhibitor LY294002. Inhibitor concentrations were chosen to be approximtaly in the mid-range of the dose-response curves for each cell line after 24 hrs of exposure (Fig. 8C). In both cell lines, inhibition of either the AR or MAPK pathways promoted activation of the PI3K pathway, as seen by the increase in phosphorylated AKT (S473) (Fig. 8A). The addition of the PI3K inhibitor, LY294002, alone or in combination diminished PI3K activity (Fig. 8A). Interestingly, the inhibition of PI3K alone, increased AR expression in both LNCaP and C4-2 cell lines (Fig. 8A). Since AR transcriptionally regulates its own expression, this suggested PI3K inhibition increased AR activity. The ribosomal protein pS6 was completely inhibited only in the presence of the PI3K inhibitor LY294002. The abundance of cleaved PARP (c-PARP), an indicator for apoptosis, was highest in the triple inhibition case for both LNCaP and C4-2 cell lines, however c-PARP also increased in the dual inhibition of RAF and PI3K in both cell lines and in the dual inhibition of PI3K and AR in C4-2 cells (Fig. 8A). We further characterized cellular viability using the MTT assay. Cell viability decreased at 72 hrs in the dual and triple inhibition cases for both LNCaP and C4-2 cell lines (Fig. 8B). However, MDV3100 (10 *μ*M) alone had only a modest effect on cell viability versus control (DMSO).

**Fig. 8:**
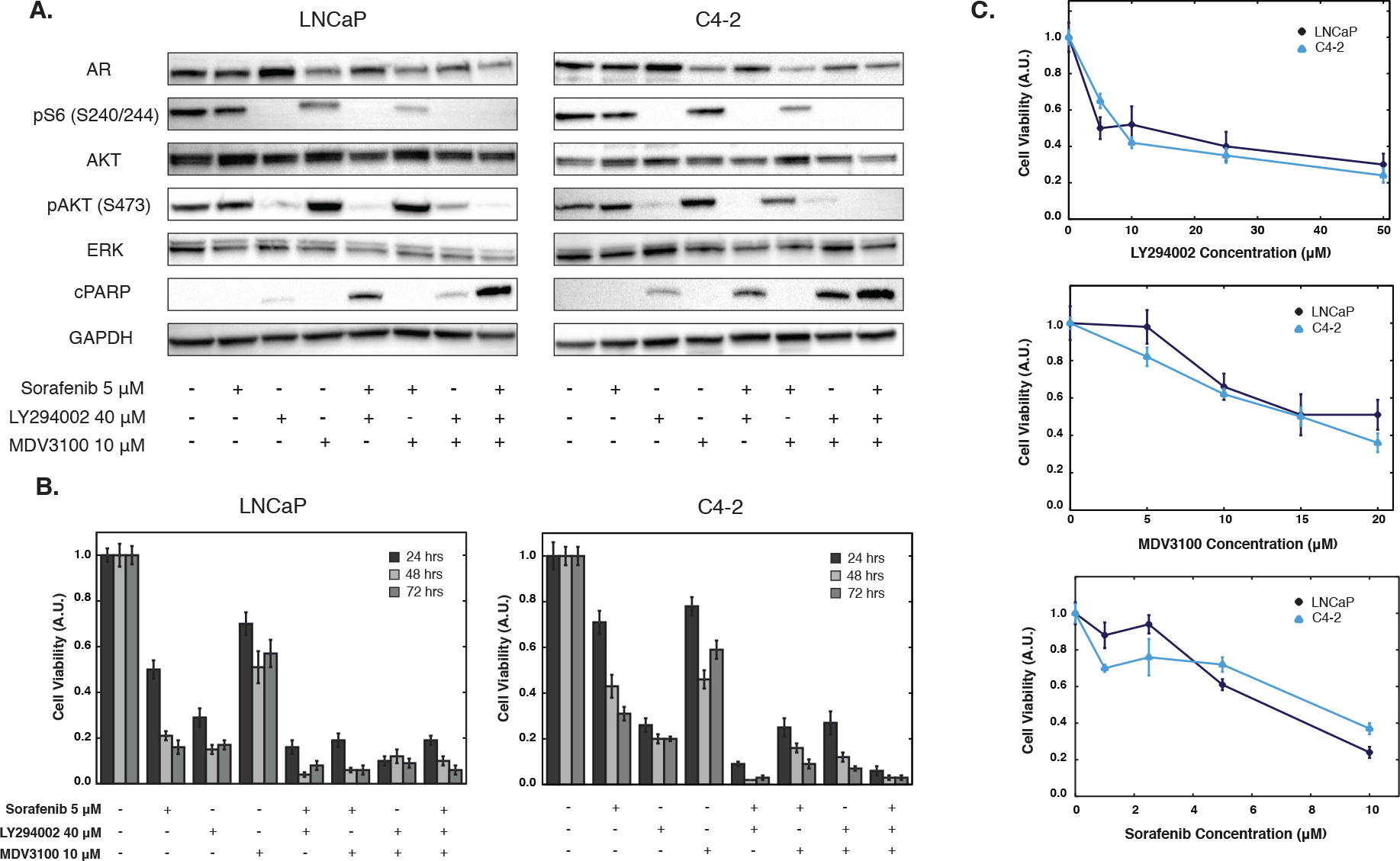
Experimental results for multiple drug combinations on two prostate cancer cell lines, LNCaP and C4-2. A. Western blot analysis of AR, pS6, AKT, pAKT, ERK and cleaved PARP in LNCaP and C4-2 cell lines treated for 24 hrs with DMSO (control), sorafenib (5 μM), LY294002 (40 μM), and MDV3100 (10 μM) alone or in combination (at least 3 repeats). B. Cells (LNCaP and C4-2) were treated for 24, 48 and 72 hrs with sorafenib (5 μM), LY294002 (40 μM), and MDV3100 (10 μM) and cell viability was measured using MTT Assay. Values were normalized to DMSO (control). C. Cell viability results for LNCaP and C4-2 cells at varying concentration of sorafenib, LY294002, and MDV3100 after 24 hrs of treatment. Values were normalized to DMSO (control). Error bars represent standard error (at least 3 repeats with triplicates performed in each experiment).

## Discussion

In this study, we analyzed a population of mathematical models that described androgen and mitogenic signaling in androgen dependent and independent prostate cancer. An ensemble of model parameters was estimated using 43 steady-state and dynamic data sets taken from androgen dependent, intermediate and independent LNCaP cell lines using multiobjective optimization. The model ensemble captured 85% of the training data, compared with 20% for the random parameter control. We tested the predictive power of the model ensemble by comparing simulations with 33 data sets (including four *in vivo* drug studies) not used for training. The model ensemble correctly predicted approximately 55% of the validation cases overall, but 75% of the clinical cases. During ensemble generation we identified potentially important missing biology. Addition of one such case, EGF-induced HER2/EGFR heterodimerization, improved both training and validation performance with no additional parameter fitting. Analysis of the model population suggested that simultaneously targeting the PI3K and MAPK pathways in addition to anti-androgen therapies could be an effective treatment for CRPC. We tested this hypothesis in both ADPC LNCaP cell lines and LNCaP derived CRPC C4-2 cells using three inhibitors: the androgen receptor inhibitor MDV3100 (enzalutamide), the Raf kinase inhibitor sorafenib, and the PI3K inhibitor LY294002. Consistent with model predictions, cell viability decreased at 72 hrs in the dual and triple inhibition cases in both the LNCaP and C4-2 cell lines, compared to treatment with any single inhibitor alone. Thus, crosstalk between the androgen and mitogenic signaling axes led to the robustness of CRPC to any single inhibitor. However, model analysis predicted efficacious target combinations which were confirmed by experimental studies in multiple cell lines, thereby illustrating the potentially important role that mathematical modeling can play in cancer.

Three of the validation cases missed by the model involved the effect of EGF on AR and AR-activated gene expression. However, the inhibition of AR activation by EGF remains an open question, with many groups debating the biology involved, particularly the role of the PI3K/AKT pathway. Multiple groups have shown decreased expression of AR and androgen-regulated PSA due to EGF stimulus in some prostate cell lines [8, 15]. Simulations of the model ensemble showed either the opposite trend or no effect due to EGF stimulus. This response may be dependent on androgen status. Lin *et al.* found that in low passage number LNCaP cells (C-33), AKT negatively regulated AR by destabilizing it and promoting ubiquitylation. On the other hand, in high passage number LNCaP cells (C-81), AKT levels were high which contributed to AR stability and less degradation [51]. Cai *et al.* found that AR protein levels in CR cells were not affected by EGF [8]. Others though have found that PSA expression, even in C-81 cells, is decreased by EGF [32]. In other prostate cell lines, EGF has been shown to increase AR transactivation [28, 68]. Much of the debated biology involves the effect of AKT activation on AR. For example, Wen *et al.* showed that HER2 induced AKT activation and LNCaP cell growth in the presence and absence of androgen [92]. While another study showed AKT phosphorylation of AR at S213 and S790 suppressed AR transactivation and AR-mediated apoptosis of LNCaP [52]. The MAPK pathway, which is downstream of EGFR, may also enhance AR responses to low levels of androgen [30, 91]. Thus, due to the discrepancies in the literature, additional experiments should be performed before revising the network connectivity to the model.

Analysis of the population of PCa models identified key signaling components and processes in AD and CR cells. There was little difference between sensitive and robust processes in AD versus CR cells in the presence of androgen. The MAPK and PI3K pathways were consistently ranked in the top 2% of sensitive species in the presence of androgen, while cell cycle species, such as cyclin D-CDK4/6 complexes bound to cell cycle inhibitors (p27Kip1, p21Cip1, p16INK4), were consistently robust. However, this profile changed considerably in the absence of androgen. The activation of PI3K and ERK by HER2 dimerization and autophosphorylation, and ERK-mediated AR activation was significantly more important in CR versus AD cells. On the other hand, although AR-regulated gene expression was equally sensitive between the cell types, transcriptional and translational processes were more robust in CR versus AD cells. This evidence supports the current theory that CR cells will still respond to androgen, and that AR can be activated in the absence of androgens by MAPK activation [22]. Thus, AR is still an active therapeutic target against CRPC [45]. Interestingly, the androgen inhibitor enzalutamide had no effect on the top 2% of sensitive species. Species in the PI3K/AKT and MAPK pathways in the presence of enzalutamide were still highly sensitive. The application of enzalutamide increased sensitivity of AR species found outside of the nucleus as well as PAcP species. Informed by our sensitivity results, we performed robustness analysis to determine the effect of combination treatments on key model proteins. Robustness analysis indicated diverse effects of Raf knock-out on PSA and cyclin D concentrations. Clinical studies of sorafenib, a multi-kinase inhibitor that has activity against Raf, showed increase PSA levels in patients [17]. Our results indicate that cell-to cell heterogeneity in gene expression can play a significant role in determining cell response. Thus, combination therapies need to be considered even in the case of a Raf knock-out.

Analysis of the population of PCa models suggested inhibition of either the PI3K or the MAPK pathways in combination with AR inhibition was a possible therapeutic strategy to treat CRPC. Carver *et al.* looked at dual inhibition of AR and PI3K signaling in LNCaP cells and in a PTEN-deficient murine prostate cancer model [64]. They found that a combination of the PI3K inhibitor, BEZ235, and the AR inhibitor, MDV3100 (enzalutamide) dramatically reduced the total cell number and increased c-PARP in the dual inhibition case. These findings lead to the hypothesis that AKT inhibition increased AR activity through increased HER3. On the other hand, AR inhibition increased AKT activity due to the down regulation of PHLPP, a protein phosphatase that regulates AKT. Dual and triple knock-out simulations of PI3K, AR (and RAF) showed only a slight additive effect on the cell cycle protein cyclin D. Thus, the combined decrease in cell population observed by Carver *et al.* was likely due to cell death and not cell-cycle arrest. The model also showed decreased cell cycle proteins in the PI3K knock-out as well as in the PI3K and AR dual knock-out case in some ensemble members. This was consistent with the decreased cell count in the PI3K inhibition case, which is not dependent on cell death as c-PARP levels were low. The decrease in cell cycle proteins in the model was due to decreased translation, including reduced levels of eIF4E and activated 40S ribosomal subunits. Decreased p70 (S6) activation due to PI3K inhibition was shown in both the model and by Carver *et al.* Lastly, our experimental results confirmed the Carver *et al.* study; dual inhibition of the AR and PI3K pathways decreased cell viability more than each of the individual inhibitors alone. We explored the addition of a third inhibitor, the Raf inhibitor sorafenib, and added an additional CR cell line, C4-2. There was no significant decrease in cell viability between the three dual inhibitor cases and the triple inhibition case at 74 hours. Thus, dual inhibition (PI3K/AR, AR/MAPK, or PI3K/MAPK) may be a sufficient treatment for CRPC.

The PCa signaling architecture was assembled after extensive literature review and hand curation of the biochemical interactions. However, there are a number of areas where model connectivity could be refined, e.g., the regulation of AR phosphorylation. We assumed a single canonical activating AR phosphorylation site (S515), with ERK being the major kinase and PP2A or PP1 being the major phosphatases responsible for regulating this site. MAPK activation following EGF treatment increases AR transcription and cell growth, partially through AR phosphorylation on MAPK consensus site S515 [68]. However, there are at least 13 phosphorylation sites identified on AR, with phosphorylation at six of these being androgen induced [25]. Moreover, other kinases such as AKT, protein kinase C (PKC) family members, as well as Src-family kinases can all phosphorylate AR in prostate cells [30, 68]. For example, AKT activation leads to AR phosphorylation at both S213 and S791, however, the role of these sites remains unclear [51, 52, 85, 92]. AKT effects on AR may also be passage number dependent, with AKT repressing AR transcription in low passage number cells and enhancing transcription in higher passage cells [51]. Androgen independent phosphorylation of AR by Src family kinases (not currently in the model) at Y534 [30] or by protein kinase C (PKC) family members at the consensus site S578 could also be important for understanding the regulation of AR activity. A second area we will revisit is the gene expression program associated with androgen action, and particularly the role of AR coregulators. Currently, we included only two AR coactivators, cyclin E and CDK6 [50, 95] and three corepressors AP1, Cdc25A, and cyclin D1a in the model [13, 67, 73]. However, there are at least 169 proteins classified as potential AR coregulators [35, 36] with many of these being differentially expressed in malignant cells. For example, the expression of steroid receptor coactivator-1 (Src-1) and transcriptional intermediary factor 2 (Tif-2), both members of the steroid receptor coactivator family, are elevated in prostate cancer [28, 29]. Src-1 is phosphorylated by MAPK and interacts directly with AR to enhance AR-mediated transcription [35]. Another class of potentially important AR coregulators are the cell cycle proteins Cdc25 and Rb. Unlike Cdc25A, Cdc25B (not in the model) can act as an AR coactivator leading to enhanced AR transcription activity [63]. The Rb protein, in addition to being a key cell cycle regulator, has been shown to be an AR coactivator in an androgen-independent manner in DU145 cells [97]. However, there is some uncertainty about the role of Rb as Sharma *et al*. showed that Rb decreased AR activation in multiple prostate cancer cell lines and xenografts [76]. Forkhead proteins have also been shown to activate as well as repress AR function. In prostate cancer, AKT suppresses AFX/Forkhead proteins, which diminishes expression of AFX target genes, such as p27Kip1 [6, 27, 59, 83]. Lastly, undoubtedly there are several other signaling axes important in PCa, such as cytokine or insulin-and insulin-like growth factor signaling [9, 37, 75, 84]. Understanding the pathways associated with these signals and how they relate to the current model, may give us a more complete picture of androgen sensitivity and progression of prostate cancer.

## Materials and Methods

Prostate model signaling architecture. We modeled the transcription, translation and post-translational modifications of key components of the PCa signaling architecture. The model, which consisted of 780 protein, lipid or mRNA species interconnected by 1674 interactions, was a significant extension to our previous model [87] in several important areas. First, we included well-mixed nuclear, cytosolic, membrane and extracellular compartments (including transfer terms between compartments). Next, we expanded the description of growth factor receptor signaling, considering both homo- and heterodimer formation between ErbB family members and the role of cellular and secreted prostatic acid phosphatase (cPAcP and sPAcP, respectively). Both forms of PAcP were included because cPAcP downregulates HER2 activity, while sPAcP promotes modest HER2 activation [90]. Third, we expanded the description of the G1/S transition of the cell cycle (restriction point). The previous model used the abundance of cyclin D as a proliferation marker, but did not include other proteins or interactions potentially important to the restriction point. Toward this shortcoming, we included cyclin E expression (and its role as a coregulator of androgen receptor expression), enhanced the description of cyclin D expression and the alternative splicing of cyclin D mRNA (including the role of the splice variants in androgen action), included the Rb/E2F pathway as well as E2F inhibition of androgen receptor expression [18], and the cyclin-dependent kinases cyclin-dependent kinase 4 (CDK4) and cyclin-dependent kinase 6 (CDK6). We also included key inhibitors of the restriction point including cyclin-dependent kinase inhibitor 1 (p21Cip1), cyclin-dependent kinase inhibitor 1B (p27Kip1), and cyclin-dependent kinase inhibitor 2A (p16INK4) [77]. Fourth, we enhanced the description of growth factor induced translation initiation. One of the key findings of the previous model was that growth factor induced translation initiation was globally sensitive (important in both androgen dependent and independent conditions). However, the description of this important subsystem was simplified in the previous model. Here, we expanded this subsystem, using connectivity similar to previous study of Lequieu *et al.* [49], and re-examined the importance of key components of this axis, such as mammalian target of rapamycin (mTOR), phosphatidylinositide 3-kinase (PI3K) and AKT. Lastly, we significantly expanded the description of the role of androgen receptor. The previous model assumed constant AR expression, consistent with studies in androgen dependent and independent LNCaP sublines [48]. However, other prostate cancer cell lines vary in their AR expression [80]. Thus, to capture androgen signaling in a variety of prostate cancer cells, we included the transcriptional regulation governing androgen receptor expression, updated our description of the regulation of androgen receptor activity and androgen action (gene expression program driven by activated androgen receptor). At the expression level, we included AR auto-regulation in combination with the co-activators cyclin E and CDK6 [50, 95]. We also assumed androgen receptor could be activated through androgen binding or a ligand-independent, MAPK-driven mechanism referred to as the outlaw pathway [22, 96]. We assumed a single canonical activating AR phosphorylation site (S515), with phosphorylated extracellular-signal-regulated kinase 1/2 (ppERK1/2) being the major kinase and protein phosphatase 2 (PP2A) or phosphoprotein phosphatase 1 (PP1) being the major phosphatases responsible for regulating this site. Finally, we modeled androgen receptor induced gene expression, including prostate specific antigen (PSA), cPAcP and p21Cip1.

**Formulation and solution of the model equations**. The prostate model was formulated as a coupled set of non-linear ordinary differential equations (ODEs):

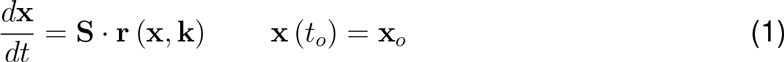

The quantity X denotes the vector describing the abundance of protein, mRNA, and other species in the model (780 × 1). The stoichiometric matrix S encodes the signaling architecture considered in the model (780 × 1674). Each row of S describes a signaling component while each column describes a particular interaction. The (*i*, *j*) element of S, denoted by *σ*_*ij*_, describes how species *i* is involved with interaction *j*. If *σ*_*ij*_ > 0, species *i* is produced by interaction *j*. Conversely, If *σ*_*jj*_ < 0, then species *i* is consumed in interaction *j*. Lastly, if *σ*_*ij*_ = 0, then species *i* is not involved in interaction *j*. The term r (x, k) denotes the vector of interactions rates (1674 × 1). Gene expression and translation processes as well as all biochemical transformations were decomposed into simple elementary steps, where all reversible interactions were split into two irreversible steps (supplemental materials). We modeled each network interaction using elementary rate laws where all reversible interactions were split into two irreversible steps. Thus, the rate expression for interaction *q* was given by:

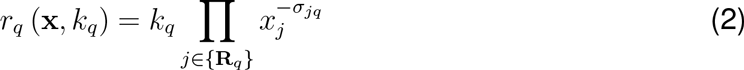

The set {**R**_*q*_} denotes reactants for reaction *q*, while *σ_jq_* denotes the stoichiometric coefficient (element of the matrix S) governing species *j* in reaction *q*. The quantity *k_q_* denotes the rate constant (unknown) governing reaction q. Model equations were generated in the C-programming language using the UNIVERSAL code generator, starting from an text-based input file (available in supplemental materials). UNIVERSAL, an open source Objective-C/Java code generator, is freely available as a Google Code project (http://code.google.com/p/universal-code-generator/). Model equations were solved using the CVODE solver in the SUNDIALS library [39] on an Apple workstation (Apple, Cupertino, CA; OS X v10.6.8).

We ran the model to steady-state before calculating the response to DHT or growth factor inputs. The steady-state was estimated numerically by repeatedly solving the model equations and estimating the difference between subsequent time points:

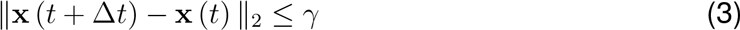

The quantities x (*t*) and x (*t* + *Δt*) denote the simulated abundance vector at time *t* and *t* + *Δt*, respectively. The *L*_2_ vector-norm was used as the distance metric, where *Δt* = 100 hr of simulated time and *γ* = 0.001 for all simulations.

We estimated an ensemble of model parameter sets using the Pareto Optimal Ensemble Techniques (POETs) multiobjective optimization routine [49, 81, 82]. POETs minimized the residual between model simulations and 43 separate training objectives taken from protein and mRNA signaling data generated in androgen dependent, intermediate and independent LNCaP cell lines (Table T1). From these training objectives, POETs generated > 10^6^ candidate parameter vectors from which we selected N = 5000 Pareto rank-zero vectors for further analysis. The set-to-set correlation between selected sets was approximately 0.60, suggesting only modest similarity between ensemble members. Approximately 33%, or 560 of the 1674 parameters had a coefficient of variation (CV) of less than 1.0, where the CV ranged from 0.59 to 5.8 over the ensemble (N = 5000). Details of the parameter estimation problem and POETs are given in the supplemental materials.

**Sensitivity and robustness analysis**. Steady-state sensitivity coefficients were calculated for N = 500 parameter sets selected from the ensemble by solving the augmented kinetic-sensitivity equations [20]:

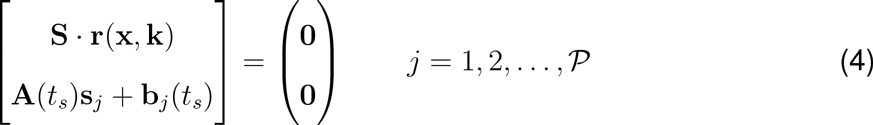

where

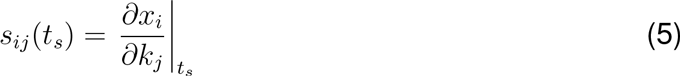
for each parameter set. Steady-state was calculated as described previously. The quantity *j* denotes the parameter index, A denotes the Jacobian matrix, and *P* denotes the number of parameters in the model. The vector b_*j*_ denotes the *j*th column of the matrix of first-derivatives of the mass balances with respect to the parameters. Steady-state sensitivity coefficients were used because of the computational burden associated with sampling several hundred parameters sets for each of the 1674 parameters. The steady-state sensitivity coefficients 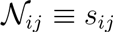 were organized into an array for each parameter set in the ensemble:

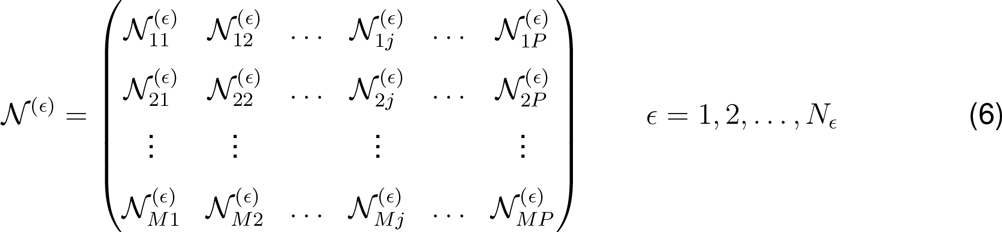

where ϵ denotes the index of the ensemble member, *P* denotes the number of parameters, *N*_*ϵ*_ denotes the number of parameter sets sampled (N = 500) and *M* denotes the number of model species. To estimate the relative fragility or robustness of species and reactions in the network, we decomposed 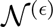 using Singular Value Decomposition (SVD):

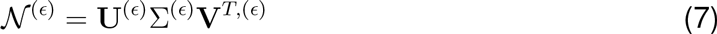

Coefficients of the left singular vectors corresponding to largest *θ* ≤ 15 singular values of 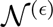 were rank-ordered to estimate important species combinations, while coefficients of the right singular vectors were used to rank important reaction combinations. Only coefficients with magnitude greater than a threshold (δ = 0.001) were considered. The fractionof the *θ* vectors in which a reaction or species index occurred was used to quantify its importance (sensitivity ranking). We compared the sensitivity ranking between different conditions to understand how control in the network shifted in different cellular environments.

Robustness coefficients were calculated as shown previously [88]. Robustness coefficients denoted by *α* (*i*, *j*, *t*_*o*_, *t*_*f*_) are defined as:

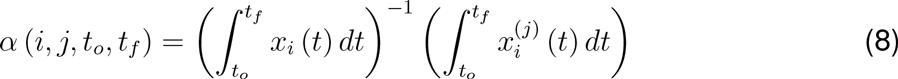

Robustness coefficients quantify the response of a marker to a structural or operational perturbation to the network architecture. Here *t*_*o*_ and *t*_*f*_ denote the initial and final simulation time respectively, while *i* and *j* denote the indices for the marker and the perturbation respectively. A value of *α* (*i*, *j*, *t*_*o*_, *t*_*f*_) > 1, indicates increased marker abundance, while *α* (*i*, *j*, *t*_*o*_, *t*_*f*_) < 1 indicates decreased marker abundance following perturbation j. If *α* (*i*, *j*, *t*_*o*_, *t*_*f*_) ~ 1 the *j*th perturbation does not influence the abundance of marker *i*. Robustness coefficients were calculated (starting from steady-state) from *t*_*o*_ = 0 hr to *t*_*f*_ = 72 hr following the addition of 10nM DHT at *t*_*o*_. For scaled log fold change we used the following equation:

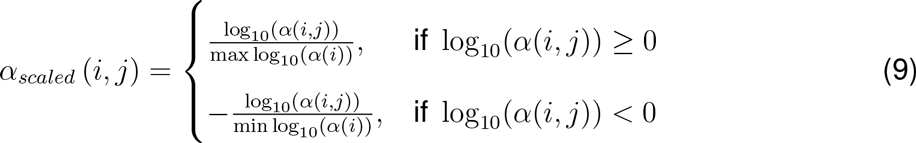

A value of *α*_*scaled*_ (*i*, *j*) > 0, indicates increased marker abundance, while *α*_*scaled*_ (*i*, *j*) < 0 indicates decreased marker abundance following perturbation *j*. If *α*_*scaled*_ (*i*, *j*) ~ 0 the *j*th perturbation does not influence the abundance of marker *i*. A value of *α*_*scaled*_ (*i*, *j*) = 1, indicates max increase of marker abundance, while *α*_*scaled*_ (*i*, *j*) = –1 indicates the max decrease of marker abundance. Robustness coefficients were calculated for the same N = 500 models selected for sensitivity analysis.

**Cell culture and treatments** Androgen dependent LNCaP prostate cancer cells were a gift from Dr. Brian Kirby (Cornell University), and the castration resistant C4-2 prostate cancer cell line was purchased from MD Anderson Cancer Center, University of Texas. Cell lines were maintained in RPMI 1640 media (Life Technologies, Inc., Grand Island, NY) with 10% fetal calf serum (FBS; Hyclone) and 1x antibiotic/antimycotic (Sigma, St. Louis, MO) in a 5% CO_2_ humidified atmosphere at 37°C. The AR inhibitor MDV3100 (enzalutamide) and the Raf inhibitor sorafenib were purchased from SantaCruz Biotechnology (Santa Cruz, CA). The PI3K inhibitor LY294002 was purchased from Cell Signaling Technologies (Danvers, MA, USA). All stock solutions were diluted in DMSO and stored at -20°C (Sigma, St. Louis, MO). Stock solution concentrations for western blotting experiments were 10 mM Sorafenib, 50 mM LY294002, and 10 mM MDV3100. For the cell viability assays, stock solution concentrations were 0.5 mM Sorafenib, 4 mM LY294002, and 1 mM MDV3100.

**Protein extraction and western blot analysis** LNCaP and C4-2 cells were seeded in 60 mm dishes at a density of 4 × 10^5^. After 96 and 72 hrs, for LNCaP and C4-2 cells respectively, the media was replaced with fresh media and drug treatments were added. After 24 hours, cells were washed twice in PBS buffer, scraped in 250 *μ*L ice-cold lysis buffer (Pierce, Rockford, IL) supplemented with protease and phosphatase inhibitors (Sigma, St. Louis, MO), and lysed for 30 min on ice. Lysates were centrifuged at 13,000 rpm for 30 min at 4°C. After quantification of total protein by BCA assay, equal amounts of total protein lysates (25 *μ*g) were resolved by SDS-PAGE and transferred onto PVDF membranes. Membranes were blocked in 5% fat free milk and then probed with antibodies. The primary antibodies used for western blot analysis were pAKT Ser473, AKT, pS6 Ser240/244, pERK Thr202/Tyr204, ERK, AR, cleaved PARP, and GAPDH were from Cell Signaling Technologies (Danvers, MA, USA). For detection, enhanced chemilumines-cence ECL reagent (GE Healthcare, Pittsburgh, PA) was used and signals were visualized using the ChemiDoc XRS system (Bio-Rad).

*MTT assay* LNCaP and C4-2 cells were seeded at a density of 1×10^4^ cells per well in 96 well plates. After 48 hrs the media was refreshed and drug treatments added. Cell growth at 24, 48, and 72 hrs was determined using a 3-(4,5-dimethyl thiazol-2-yl)-2,5-diphenyl tetrazolium bromide (MTT) assay. At the specified time point 10 *μ*L MTT reagent (stock of 5 mg/mL in PBS) was added to each well and the cells were further incubated for 4 hrs. At 4 hrs, the media was removed and 50 *μ*L of dissolving reagent DMSO was added to each well. After an additional 10 min incubation, the absorbance was measured at 540 nm on a microplate reader. Each reading was adjusted by subtracting the absorbance value for the blank (media only) and the results were then scaled to the DMSO-treated (control) case.

## Acknowledgements

This study was supported by award number U54CA143876 from the National Cancer Institute. The content is solely the responsibility of the authors and does not necessarily represent the official views of the National Cancer Institute or the National Institutes of Health. The authors acknowledge the financial support of the National Science Foundation, award no. DGE-1045513,for a GK12 training grant entitled (Grass Roots: Advancing Education in Renewable Energy and Cleaner Fuels through collaborative graduate fellow/teacher/grade-school student interactions.We would like to thank Timothy Chu, Kyr-iakos Tsahalis, Spencer Davis, Rohit Yallamraju, Gongcheng Lu, and Yalin Zhao for model development, and cell culture studies.

## Supplementary materials

**Estimation of a population of models using Pareto Optimal Ensemble Techniques (POETs)**. We used multiobjective optimization to estimate an ensemble of prostate models. Although computationally more complex than single-objective formulations, multiobjective optimization can be used to address qualitative conflicts in training data arising from experimental error or cell-line artifacts [33]. In this study we used the Pareto Optimal Ensemble Technique (POETs) to perform the optimization. POETs integrates standard search strategies, e.g., Simulated Annealing (SA) or Local Pattern Search (PS) with a Pareto-rank fitness assignment [81]. The mean squared error, *η*, of parameter set k for training objective *j* was defined as:

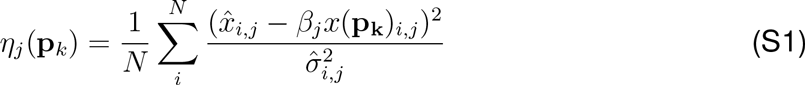

The symbol 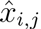 denotes scaled experimental observations (from training objective j) while *x*(p_k_)_*i*,*j*_ denotes the simulation output (from training objective j). The quantity *i* denotes the sampled time-index or condition, and *N* denotes the number of time points or conditions for experiment j. The standard deviation, 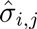, was assumed to be equal to 10% of the reported observation, if no experimental error was reported. *β*_*j*_ is a scaling factor which is required when considering experimental data that is accurate only to a multiplicative constant. In this study, the experimental data used for training and validation was typically band intensity from immunoblots, where intensity was estimated using the ImageJ software package [1]. The scaling factor used was chosen to minimize the normalized squared error [5]:

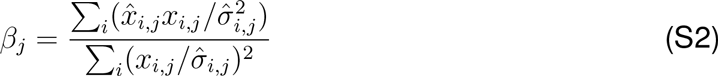

By using the scaling factor, the concentration units on simulation results were arbitrary,which was consistent with the arbitrary units on the experimental training data. All simulation data was scaled by the corresponding *μ*_*j*_.

We computed the Pareto rank of parameter set k_*i*+1_ by comparing the simulation error at iteration *i*+1 against the simulation archive, denoted as K*_i_*. We used the Fonseca and Fleming ranking scheme [23] to estimate the rank of the parameter set k_*i*+1_. Parameter sets with increasing rank are progressively further away from the optimal trade-off surface.The parameter set k_*i*+1_ was accepted or rejected by the SA with probability 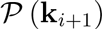:

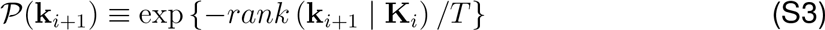

where *T* is the computational annealing temperature. The Pareto rank for k_*i*+1_ is denoted by *rank*(k_*i*+1_|K_*i*_. The annealing temperature was adjusted according to the schedule T_*k*_ = *μ*^*k*^T_*0*_ where *μ* was defined as 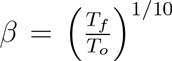. The initial temperature was given by *T*_*0*_ = *n*/*log*(2), with *n* = 4 and the final temperature *T*_*f*_ = 0.1 used in this study. The epoch-counter *k* was incremented after the addition of 50 members to the ensemble. As the ensemble grew, the likelihood of accepting a high rank set decreased. Parameter sets were generated by applying a random perturbation in log space:

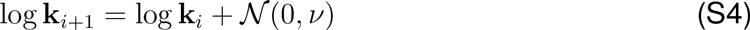

where 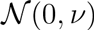 is a normally distributed random number with zero mean and variance *v*,set as 0.1 in this model. The perturbation was applied in log space to account for large variation in parameter scales and to ensure positive parameter values. We used a local pattern search every *q* steps, in our case 20, to minimize error for a single randomly selected objective. The local pattern-search algorithm used has been described previously [24].

**Translation and Transcription Template** We utilized the following template for the transcription of genes in the network without a transcription factor:

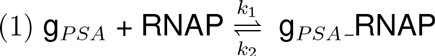

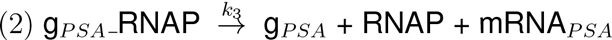

and with a transcription factor:

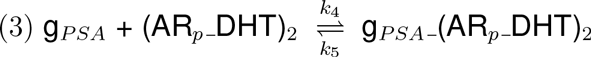

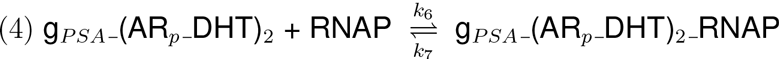

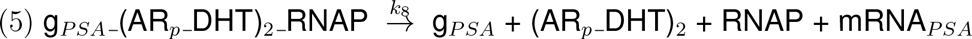

RNAP denotes RNA polymerase. Next translation was modeled by the following, where Ribo denotes ribosome:

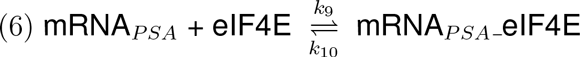

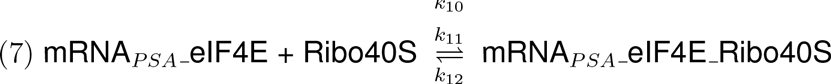

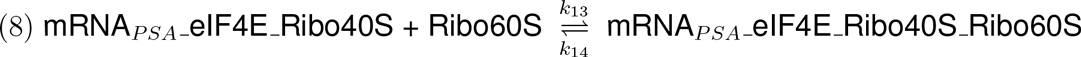

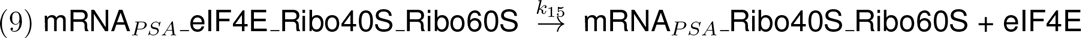

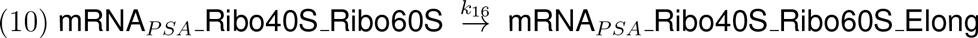

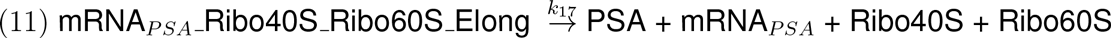

**Fig. S1:**
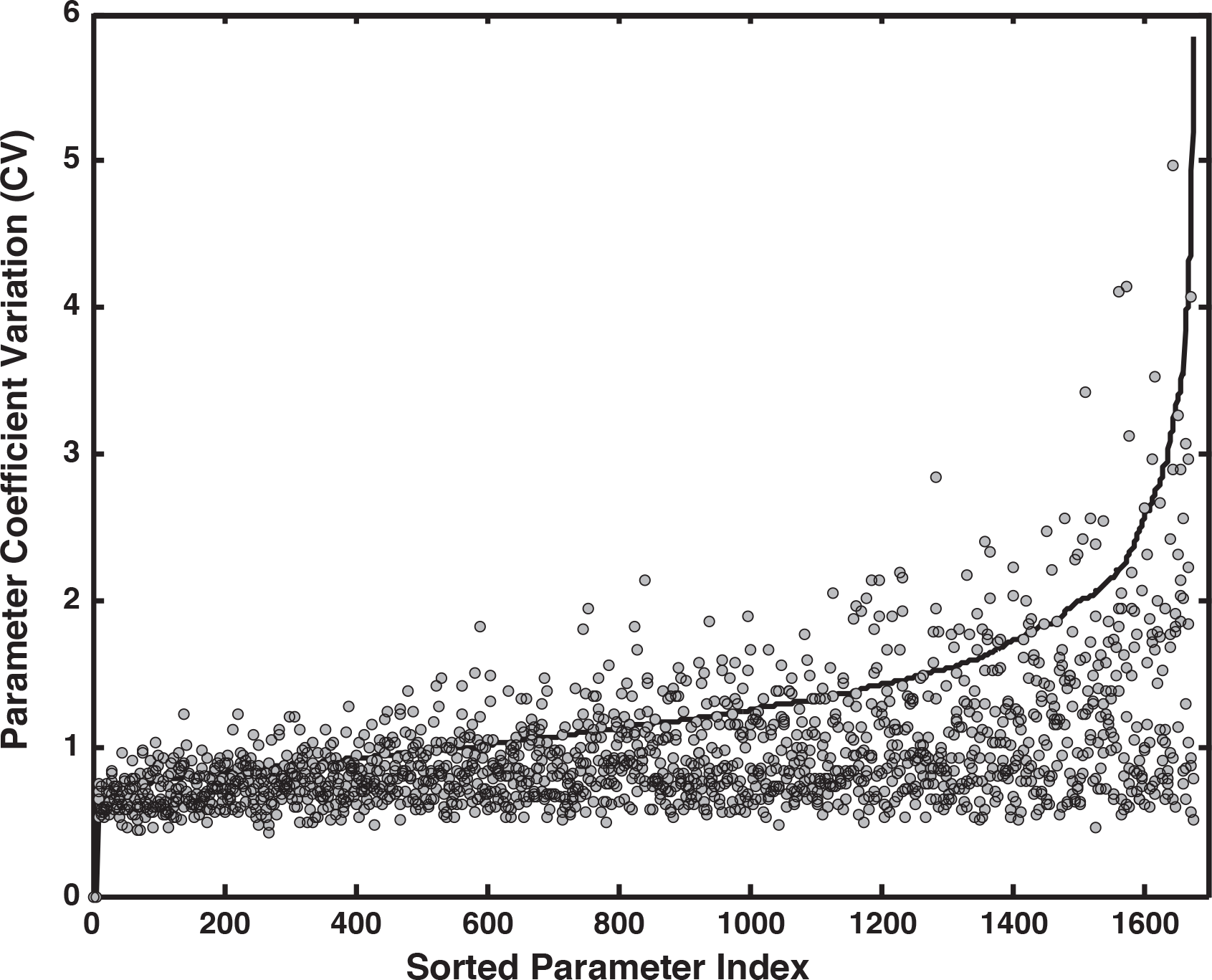
Coefficient of variation (CV) of model parameters estimated using POETs. The solid line denotes the mean CV calculated over the entire ensemble (N = 5000). The points denote the mean CV of the 500 ensemble members used for sensitivity and robustness calculations. Over the ensemble, the coefficient of variation (CV) of the kinetic parameters spanned 0.59 – 5.8, with 33% of the parameters having a CV of less than one.

**Fig. S2:**
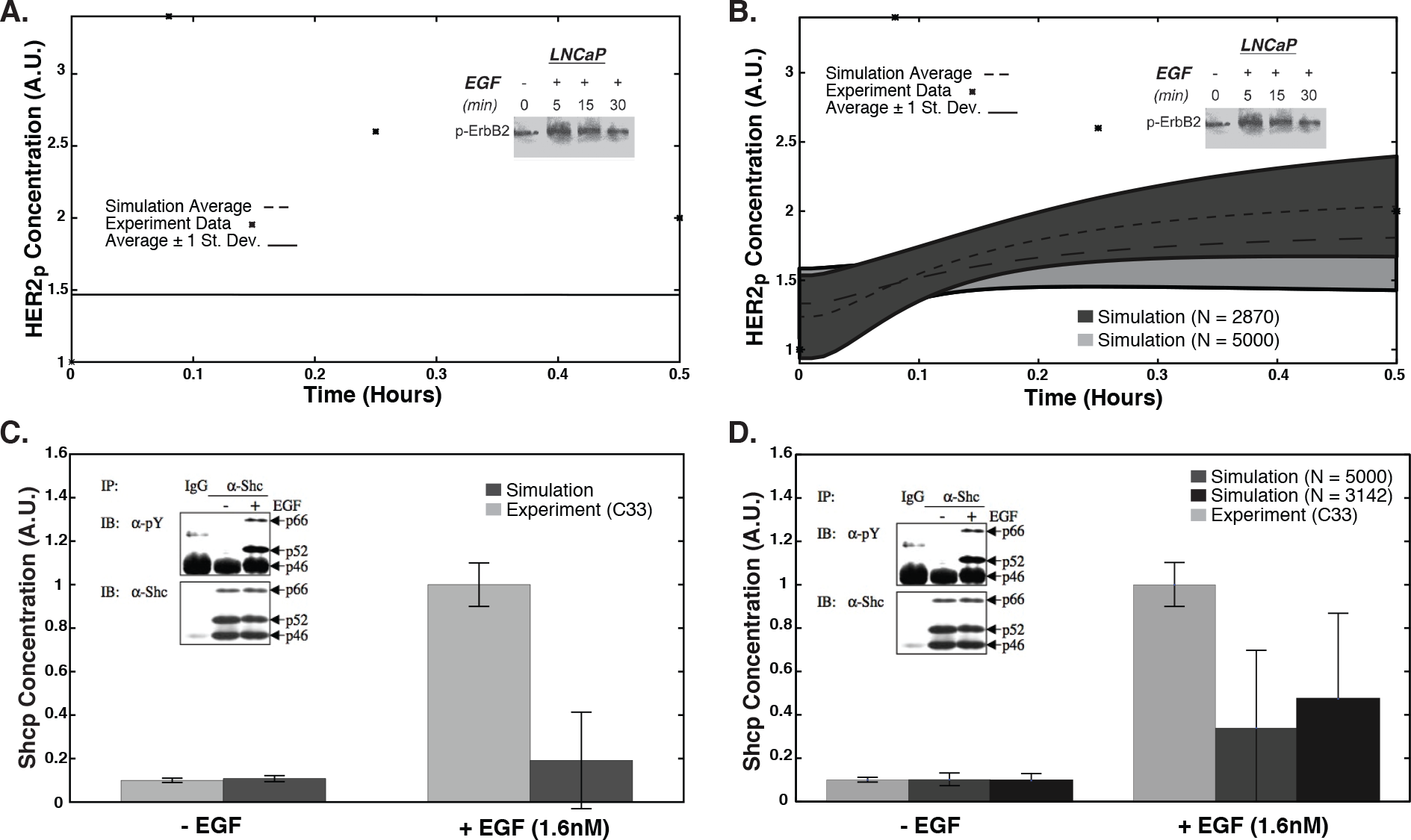
Blind model predictions for the ensemble with the original and updated model (EGFR and HER2 heterodimer). A,B. Time course data for HER2 phosphorylation due to a stimulus of 1.6 nM EGF (LNCaP C33, P7) for the old and new model, respectively. Dark grey shows only parameters improved by the updated model (N=2870) while light grey show all parameter sets (N=5000). C,D. Shc phosphorylation levels at 16 hrs in the presence and absence of 1.6 nM EGF (LNCaP C33, P29) for the old and new model, respectively. Light grey denotes experimental data, mid grey denotes simulation results for all parameters (N=5000), and black denotes only parameters improved by the updated model (N=3142). Error bars denote plus and minus one standard deviation from the mean.

**Fig. S3:**
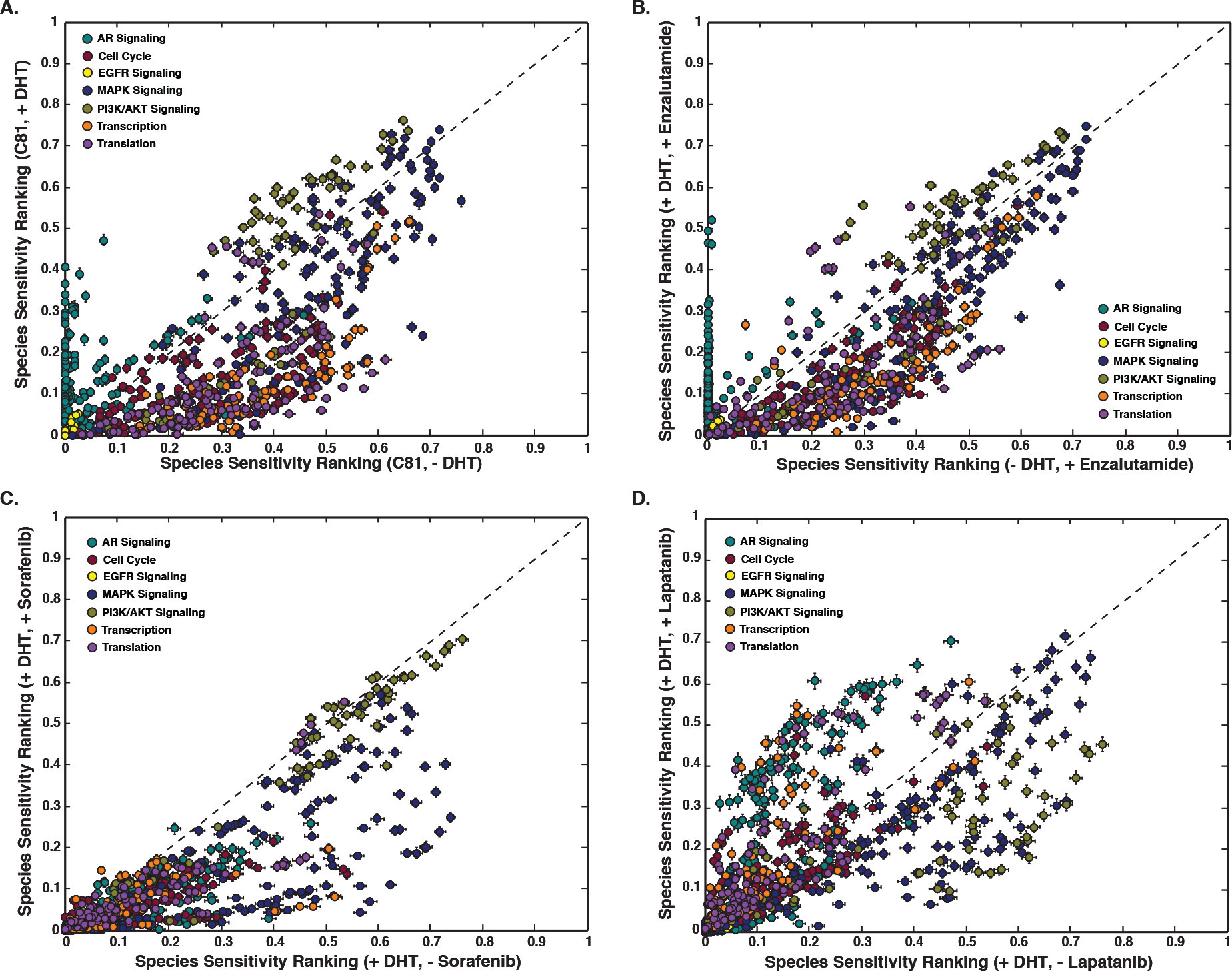
Sensitivity analysis of a population of prostate models (N = 500). Species with a low sensitivity are considered robust, while species with a high sensitivity ranking are considered fragile. A Sensitivity ranking of network species in CR cells in the absence and presence of DHT. B. Sensitivity ranking of network species in CR cells in the presence of enzalutamide in the presence and absence of a DHT stimulus. C., D. Sensitivity ranking of network species in CR cells in the presence and absence of sorafenib and lapatinib, respectively, with a DHT stimulus. Error bars denote standard error with N = 500.

**Fig. S4:**
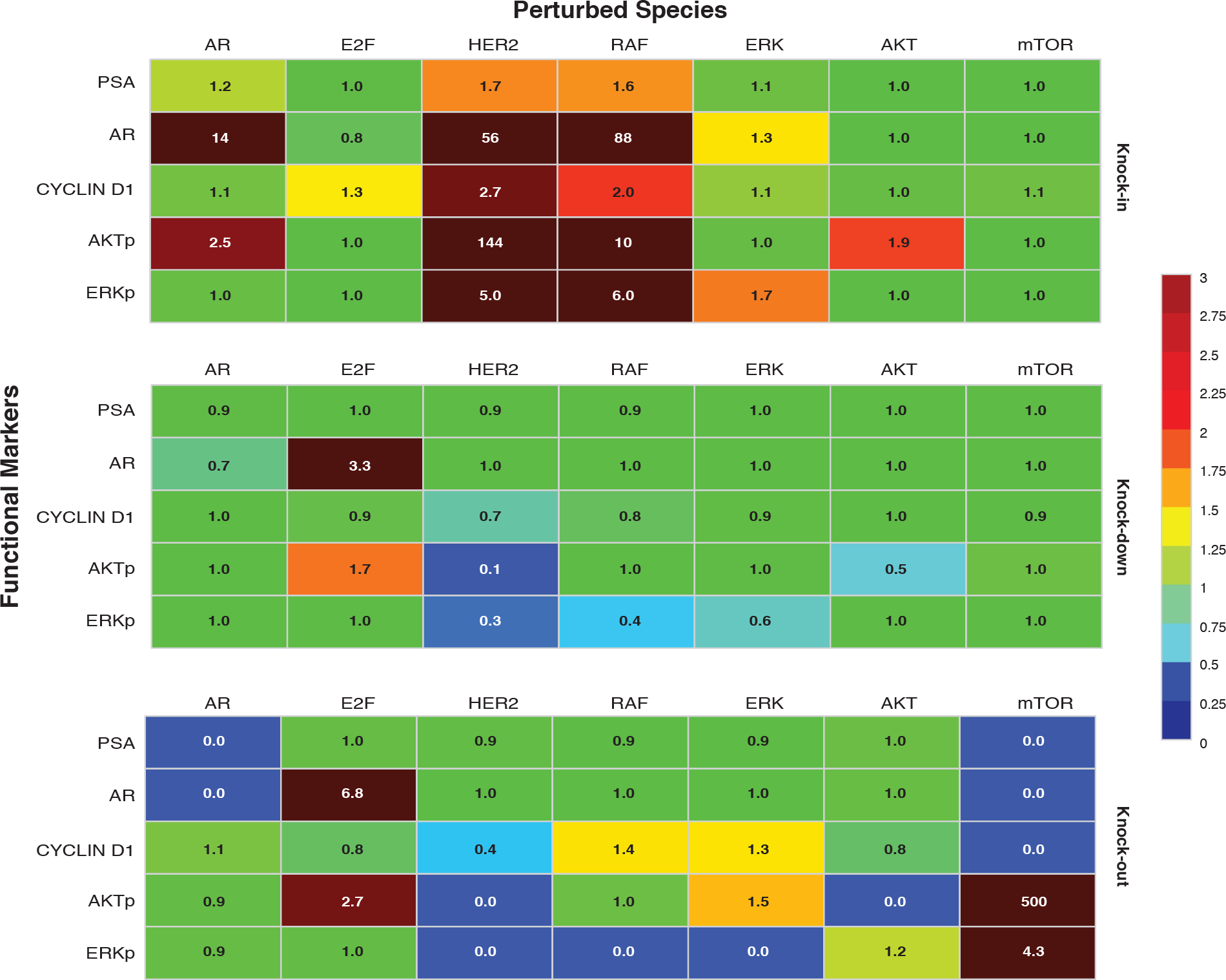
Robustness analysis of protein markers. Expression level of key proteins was altered by a factor of 2, 0.1, or 0 (knock-in, knock-down, or knock-out) and robustness coefficients were calculated for five key protein markers. Simulations shown were from CR cells, with indicated perturbation. Mean of 500 ensemble members is shown.

**Fig. S5:**
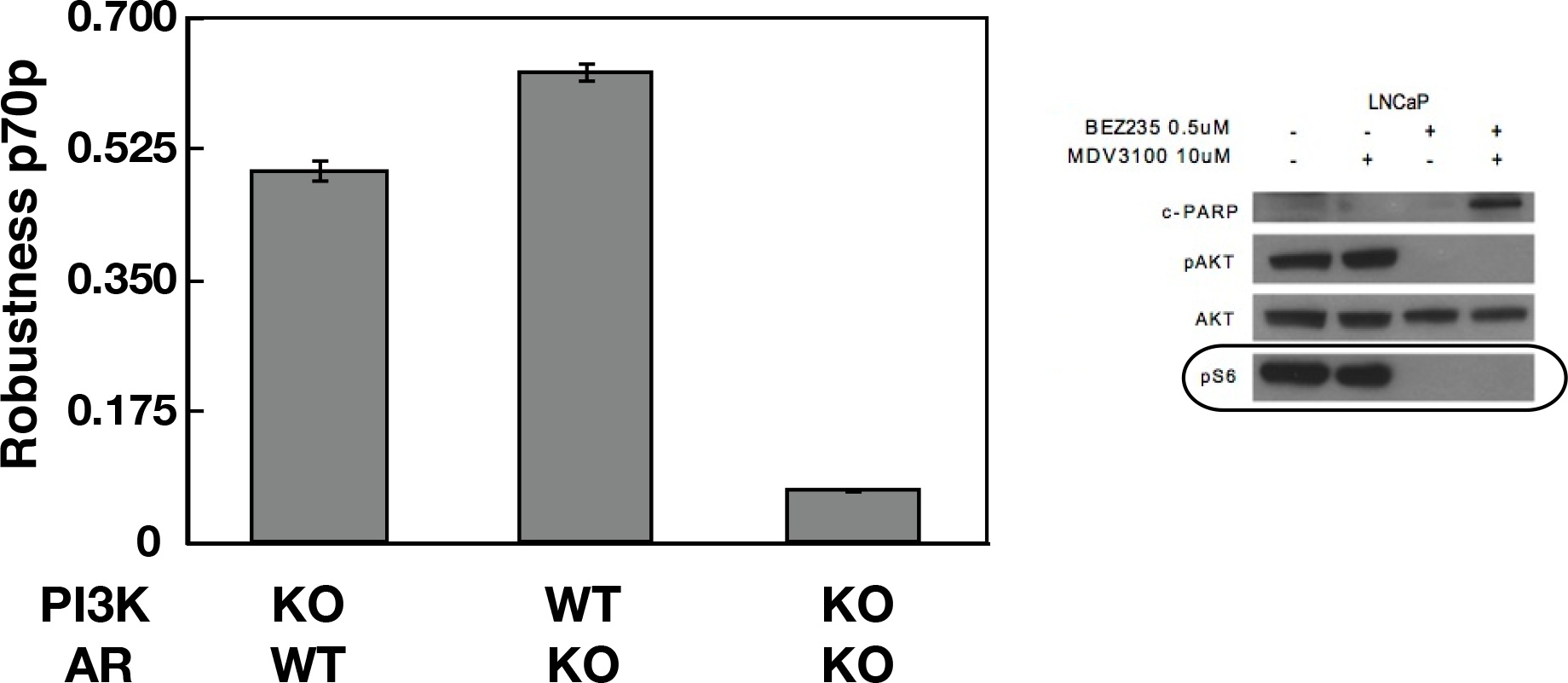
Robustness analysis of protein markers. Expression level of key proteins was altered by a factor of 2, 0.1, or 0 (knock-in, knock-down, or knock-out) and robustness coefficients were calculated for five key protein markers. Simulations shown were from CR cells, with indicated perturbation. Mean of 500 ensemble members is shown.

